# Acute Hypoxia Disrupts Multiple Processes Supporting Spatial Navigation

**DOI:** 10.64898/2026.09.24.753962

**Authors:** Daniel J McKeown, Heiki M Bahnson-Wright, Trinity Caldwell, Madeliene Marks, Georgia R Marshall, Holly Newton, Dylan Stenholm, Michael Young, Douglas J Angus, Victor R Schinazi

## Abstract

Acute hypoxia impairs cognitive function, yet its effects on navigation and intrinsic brain activity remain poorly understood. This study investigated whether acute continuous hypoxia (ACH) and intermittent hypoxia (AIH) differentially influence navigation performance and resting-state electroencephalography (EEG) in healthy young adults. Seventy-two participants completed sham (FiO□ = .209) and hypoxia (FiO□ = .13) sessions during either a 1-hour ACH or AIH intervention. Navigation was assessed using the Spatial Performance Assessment for Cognitive Evaluation (SPACE), while resting-state EEG was recorded before and after the intervention in a subset. EEG spectra were parameterised into periodic (alpha and beta power and central frequency) and aperiodic (exponent and offset) components. Hypoxia impaired multiple components of navigation, increasing path integration distance error and egocentric pointing angle error irrespective of intervention group membership, while mapping accuracy was preferentially impaired following AIH and perspective taking performance following ACH. Hypoxia also reduced alpha and beta power, beta central frequency, and aperiodic exponent, while aperiodic offset and alpha central frequency remained stable. The reduction in aperiodic exponent was primarily evident following AIH, suggesting that the temporal pattern of oxygen delivery may influence specific neurophysiological responses. SpO□ reduction did not predict individual differences in navigation performance or EEG activity. These findings demonstrate that brief, moderate hypoxic exposure can disrupt allocentric and egocentric components of spatial navigation while selectively altering periodic and aperiodic features of intrinsic brain activity.

## INTRODUCTION

Each year, millions of people worldwide visit or reside in high-altitude environments where oxygen availability is reduced. Exposure to high-altitude environments is also a regular occupational demand for aviation, defence, and first-responder personnel (Gatterer et al., 2024). High-altitude environments present both physiological and psychological challenges. When physiological adaptations fail to adequately compensate for reduced atmospheric oxygen availability, reductions in peripheral blood oxygen saturation (SpO_2_; hypoxemia) may occur, limiting oxygen delivery to tissues (Ainslie & Subudhi, 2014). If neural tissue experiences hypoxia, neurophysiological impairments may emerge, potentially disrupting complex cognitive functions (McKeown, Angus, et al., 2025; McMorris et al., 2017; Taylor et al., 2015). Among these functions, navigation may be particularly vulnerable, as it relies on the coordination of multiple brain regions involved in attention, memory, and executive control (Baumann & Mattingley, 2021; Ekstrom & Hill, 2023; R. A. Epstein et al., 2017). These metabolically demanding processes are necessary to support the flexible manipulation of spatial information and the adaptation of strategies needed to solve complex environmental challenges (Baumann & Mattingley, 2021; Burgess, 2006; Ekstrom & Hill, 2023; Epstein, 2008). Understanding how reduced oxygen availability affects the neural processes supporting navigation may therefore have important implications for both everyday functioning and occupational performance in high-altitude settings.

### Neural mechanisms of navigation

Since Tolman‘s seminal proposal that animals form cognitive maps of their environment, our understanding of navigation has evolved substantially (Epstein et al., 2017; Tolman, 1948). Tolman argued that successful navigation depends not only on learned stimulus-response associations but also on internal representations of encoded spatial relationships between locations. Such representations allow individuals to plan novel routes, take shortcuts, and adapt flexibly to changing environments. Building on this foundation, subsequent research in animals and humans has identified the distributed involvement of neural networks responsible for supporting place- and response-based navigation strategies (Baumann & Mattingley, 2021; Farzanfar et al., 2023; Iaria et al., 2003; Packard & McGaugh, 1996).

Successful place-based navigation relies on encoding landmarks and developing allocentric representations of the environment. This strategy allows flexible route planning, shortcut finding, especially in unfamiliar environments (Doeller et al., 2008; Iaria et al., 2003; Konishi & Bohbot, 2013; Wegman et al., 2014). In contrast, response-based strategies rely on “person-centred” egocentric representations and memorising sequences to tackle immediate spatial problems during navigation (i.e., ‘do I turn left or right at the fork in the road?’). Response-based strategies are effective during route learning because they rely on memorising the necessary steps to reach your destination when you are unable to develop a flexible cognitive map of the environment. However they become less effective when an individual must adapt their route due to unforeseen perturbations (Burgess, 2006; Iaria et al., 2003; Iglói et al., 2009; Packard & McGaugh, 1992).

The hippocampus plays an important role in supporting place-based navigation (Ekstrom & Ranganath, 2018; O’Keefe & Nadel, 1979; Wilson & McNaughton, 1993). Hippocampal place cells, first identified through recording in freely moving rats, activate when the animal occupies a specific location within an environment (O’Keefe & Dostrovsky, 1971). Indeed, by recording from many hippocampal neurons simultaneously, Wilson and McNaughton (1993) showed that their combined activity could be used to reconstruct the animal’s location as it moved through the environment. Evidence of comparable location coding has also been reported in humans. Single-cell recordings showed that hippocampal cells responded selectively to particular locations during virtual navigation (Ekstrom et al., 2003; O’Keefe & Dostrovsky, 1971; O’Keefe & Nadel, 1978).

Converging evidence from human structural imaging, functional, and lesion studies further corroborates these findings. Hippocampal volume has been linked to greater navigation expertise (Maguire et al., 1998, 2006), larger posterior hippocampal volume has been associated with more accurate knowledge of spatial relationships in a newly learned environment (Schinazi et al., 2013), and greater hippocampal grey matter density has been associated with the use of spatial rather than response strategies during navigation (Bohbot et al., 2007; Konishi & Bohbot, 2013). Functional imaging studies have shown that hippocampal activity increases during navigation tasks, particularly in place-based navigation, spatial memory retrieval, route planning, and proximity to a goal (Hartley et al., 2003; Spiers & Gilbert, 2015; Viard et al., 2011), while lesion studies have found that damage to the hippocampus impairs spatial learning, allocentric spatial memory, and overall navigation (Abrahams et al., 1997; Astur et al., 2002; Spiers et al., 2001).

In contrast, the striatum, particularly the caudate nucleus, plays an important role in supporting response-based navigation learning by chaining specific actions at designated locations. Evidence from rats shows that the temporary inactivation or lesion of the caudate nucleus can impair response learning, but not place learning (Packard & McGaugh, 1992, 1996). In humans, structural imaging studies have linked the use of response strategies to greater grey matter volume (Bohbot et al., 2007; Sodums & Bohbot, 2020), while functional imaging has associated activity in this region with following familiar routes (Hartley et al., 2003; Iaria et al., 2003). Critically, the hippocampal and striatal system can work together to support successful navigation (Brown et al., 2012; Doeller et al., 2008; Voermans et al., 2004).

The hippocampus and striatum are part of a broader network of regions that function together to support navigation (Baumann & Mattingley, 2021; Epstein et al., 2017). Within the entorhinal cortex, grid cells in animals (Hafting et al., 2005; Yartsev et al., 2011) and humans (Jacobs et al., 2013) fire in a regular hexagonal grid pattern across the environment. This pattern is thought to provide an internal spatial coordinate system that supports path integration, allowing animals and humans to keep track of their position and orientation during movement. Completing this representation of position, head-direction cells recorded in the postsubiculum of rats, encode facing direction irrespective of position (Taube et al., 1990a, 1990b), while the cerebellum interacts with this system and contributes to spatial updating by processing self-motion information (Cullen & Taube, 2017; Rondi-Reig et al., 2014, 2022).

In humans, navigation also draws on regions involved in scene processing, including the parahippocampal place area, the occipital place area, and the retrosplenial cortex (Epstein, 2008; Epstein et al., 2007; Epstein & Kanwisher, 1998). These regions contribute to scene recognition and visually guided navigation (Persichetti & Dilks, 2018), with the retrosplenial cortex known to support transformations between egocentric and allocentric reference frames (Marchette et al., 2014). Navigating complex environments regularly requires transitioning attention between competing demands to effectively plan routes, maintain goals, and adapt behaviours, especially when environmental restraints change (e.g., unexpected detours). These higher-order executive functions are primarily supported by the prefrontal cortex (Javadi et al., 2019; Spiers & Gilbert, 2015; Spiers & Maguire, 2006).

### Navigation in hypoxic environments

Hypoxia impairs key cognitive processes implicit in efficient navigation (Asmaro et al., 2013; McKeown, Angus, et al., 2025; McMorris et al., 2017; Taylor et al., 2015; Turner, Byblow, et al., 2015), especially memory and executive functioning. However, hypoxia’s effects on spatial navigation are less clear. Rodent models have shown that hypoxia can impair (Biswal et al., 2016; Lin et al., 2012; Zhu et al., 2019) but also improve (Wittner & Ríha, 2005) navigational ability. Similarly, research with humans in hypoxia has shown that egocentric ability and visuospatial function can be impaired (Ando et al., 2010; Barbur & and Connolly, 2011; Blacker et al., 2021; Blacker & McHail, 2022; Bondi et al., 2021; Hutcheon et al., 2023; Saputra et al., 2022) but at times improve (Schlaepfer et al., 1992). These findings suggest that hypoxia modulates the activity of neural networks responsible for key processes needed for effective navigation

The hippocampus is particularly susceptible to hypoxia (Biswal et al., 2016; Hencz et al., 2023; Lu et al., 2025; Micaux et al., 2025; Shushanyan et al., 2023; Vetrovoy et al., 2021; Zhang et al., 2018; Zhu et al., 2019). Hippocampal susceptibility results in long-term potentiation and hyperexcitability (Ai & Baker, 2006; Crepel et al., 1993; Ji et al., 2021; Kumral et al., 2004; Maiti et al., 2008; Quintana et al., 2015), disruptions to modulatory neurotransmitter activity (e.g., reduced ACh production; Bekker et al., 2006; Gibson & Duffy, 1981), and altered firing of excitatory and inhibitory neurones (Hencz et al., 2023), in CA1 and CA3 regions. Considering this, impairments in navigational strategies reliant on the hippocampus likely occur when exposed to continuous hypoxia and can consequently impair an individual’s ability to map out an environment and adopt different spatial perspectives (Chrastil et al., 2015; Lambrey et al., 2012; Wolbers et al., 2007).

Intermittent hypoxia is a therapeutic strategy involving repeated cycles of brief exposure to low oxygen followed by a return to normal oxygen levels (Burtscher et al., 2024; Navarrete-Opazo & Mitchell, 2014). While showing promise as an effective therapy for improving exercise capacity and neuronal function (Christiansen et al., 2018, 2021; Navarrete-Opazo & Mitchell, 2014; Ruggiero et al., 2020), it’s effectiveness at improving cognitive function is mixed (Bayer et al., 2017; Boulares et al., 2024; Burtscher et al., 2024; Champod et al., 2013; Navarrete-Opazo & Mitchell, 2014; Wang et al., 2020). Animal models have demonstrated that hippocampal-dependent navigational tasks are vulnerable to intermittent exposures of hypoxia (Serebrovska et al., 2025; Zhang et al., 2006). Intermittent hypoxia has been demonstrated to improve executive function in older adults (Bayer et al., 2017; Schega et al., 2016; Wang et al., 2020) but impair working memory in young adults (Champod et al., 2013). This suggests that the impact of intermittent hypoxia may elicit task-, dose-, and age-specific neuroprotective and cognitive-enhancing effects (Burtscher et al., 2024; Navarrete-Opazo & Mitchell, 2014). Possible mechanisms include increased stabilisation of hypoxia-inducible factors (HIF-α), changes in metabolic signaling, and activation of antioxidant pathways (Burtscher et al., 2024). While much of this work has been done in elderly populations where general cognitive decline has occurred with age (Bayer et al., 2017; Schega et al., 2013, 2016; Wang et al., 2020), little is known on how this translates to younger adults and during tasks that require greater cognitive demand (i.e., navigating an environment). Whether the presence of improved or impaired navigational ability occurs in a healthy young population, where cognitive function is assumably in-tact, has yet to be explored.

### EEG correlates of hypoxia

In environmental exposures to hypoxia, changes in EEG activity are inconclusive and appear to depend on the delivery type, severity, and duration of exposure (Nakata et al., 2025; Otto et al., 2025). Indeed, delta, alpha, and beta power have been reported to increase but also decrease when exposed to hypoxia (Chang et al., 2022; Feddersen et al., 2007; Hutcheon et al., 2023; Rice et al., 2019a, 2019b; Zhao et al., 2016). Although increases in alpha power have been reported to coincide with increases in adverse symptoms during moderate hypoxic exposures, reductions in alpha power that are not initially problematic at moderate hypoxic exposures have also been reported to coincide with more severe symptomatology once severe hypoxic exposures are reached (Sand & Nygaard, 1998). Similarly, increases in delta activity in severe hypoxia have coincided with ineffective physiological adaptation (i.e., hyperventilation that does not adequately ameliorate hypoxia) and greater adverse symptoms (Feddersen et al., 2007). Considering this, changes in resting-state neural dynamics may, in part, scale with the severity of exposure. EEG recorded during cognitive task provides further evidence that hypoxia modulates neural processing. Here, event-related potentials provide evidence that cortex-wide slowing in neural networks occurs with hypoxia (Blacker et al., 2021, 2021; Singh et al., 2003; Wang et al., 2014). This suggests that the impact of hypoxia coincides with both intrinsic and task-dependent changes in neural activity.

One explanation for the divergent findings in EEG measures during hypoxia is that hypoxia affects underlying cortical dynamics that are not adequately captured by conventional frequency-band measures, but may be reflected in aperiodic activity of the EEG (Preston et al., 2026). During spectral analysis of the EEG, aperiodic activity can be quantified with an exponent (the slope of the power spectral density) and offset (the y-intercept of the power spectral density) value (Donoghue et al., 2020). The aperiodic component of the EEG has recently received particular attention, and relates to pathological and non-pathological cognitive decline (Brake et al., 2024; Donoghue, 2024; Finley et al., 2024; McKeown, Roberts, et al., 2025; Voytek et al., 2015). Here, greater aperiodic activity (reflected by greater exponent and offset values) are correlated with better cognitive performance (Cazares et al., 2026; Finley et al., 2024; McKeown, Roberts, et al., 2025; Thuwal et al., 2021; Voytek et al., 2015).

In the only current assessment of aperiodic EEG activity during hypoxia, exposure to high altitude not only reduced alpha power but also reduced the aperiodic exponent, potentially reflecting increased global network excitability (Coronel-Oliveros et al., 2023). However, evidence from related respiratory manipulations suggests that aperiodic activity may not respond uniformly to changes in respiratory physiology. For example, aperiodic activity remained unchanged following experimentally induced hyperventilation, an acute respiratory response that can accompany hypoxia (Yang et al., 2025). However, these findings do not establish how intermittent hypoxia affects aperiodic EEG activity. This is an important consideration given that intermittent hypoxia may elicit a hormetic stress response, whereby brief exposures to adverse stimuli can induce adaptive changes, including enhanced synaptic plasticity (Boulares et al., 2024; Schega et al., 2016). Such adaptive responses may also influence intrinsic brain activity and could be reflected in changes in aperiodic EEG activity.

This paper reports the outcomes of two experiments, one investigating a virtual navigation test battery, the Spatial Performance Assessment for Cognitive Evaluation (SPACE), during 1 hour of acute continuous hypoxia (ACH). The second investigated SPACE performance over the same duration of hypoxia but also included an alternative acute intermittent hypoxia (AIH) delivery protocol (one minute of hypoxia, one minute of ambient air). Ten minutes of resting state EEG activity was also recorded prior to performing SPACE. As hypoxia has been linked to modulated cognitive performance and EEG activity, three hypotheses were derived. Firstly, ACH will reduce SPACE performance, whereas AIH will improve it. Second, ACH will reduce aperiodic EEG activity, while AIH will increase it once controlling for changes in periodic EEG activity. Third, individuals with greater reductions in SpO_2_ will have greater reductions in SPACE performance and aperiodic EEG activity.

## METHODS

### Ethical approval

Each participant gave written and verbal informed consent prior to undertaking testing at each laboratory visit. The study conformed to the standards set in the *Declaration of Helsinki.* The Bond University Human Research Ethics Committee approved this study (reference number: DM03041).

### Participants

Seventy-two healthy adults participated across the two experiments between February 2024 to July 2025 (18 – 37 years old, mean age = 21.78, SD = 4.95; 9 male). All participants completed a medical history questionnaire prior to their first visit which included exclusion criteria related to cardiorespiratory, neurological, and psychological conditions, smoking/vaping status, implanted medical devices, and pregnancy. Participants were required to refrain from heavy physical or cognitively demanding tasks 12 hours leading up to testing and were required to abstain from any central nervous system stimulants or depressants (e.g., caffeine and alcohol). All participants had not been exposed to a real or simulated high-altitude environment, or hypoxic stimulus within the past 6 months. For all but four participants, this was their first experience of a high-altitude environment. Statistical power was determined using G*Power (v3.1.9.4) based on an ANOVA repeated measure, within-between interaction approximation with a medium effect size (*f* = .25). To achieve a statistical power of .95 at an α of .05, a minimal sample size of 54 was sufficient. Three participants withdrew from the study due to intolerable symptoms from the hypoxic stimulus (i.e., moderate-to-severe dizziness, nausea, and headache).

### Experimental Design

The study consisted of ACH and AIH interventions. Each intervention was a two-way, sham-controlled, counterbalanced design where participants attended the laboratory on two separate occasions separated by at least 7 days to perform a hypoxia session and a sham session. This was a single-blinded study, as the investigators were required to know whether the sham or hypoxia session was being delivered to the participants for safety considerations. For the hypoxia session, the fraction of inspired oxygen (FiO_2_) was clamped at 0.13 (equivalent to ∼ 3800 - 4000 m above sea level) and was maintained for a total time of 1 hour. In the sham session, FiO_2_ remained at 0.21 (ambient air) throughout the visit. Due to the counterbalanced design of the study, half the participants were exposed to the hypoxia session on their first visit, while half of the participants were exposed to the sham session on their first visit. The ACH intervention consisted of delivering the hypoxic stimulus continuously for 1 hour following a 15-minute titration phase. During the AIH intervention, participants were exposed to 30 cycles of 1-minute hypoxic air interspersed with 1-minute of ambient air without a 15-minute titration phase.

### Instrumentation

#### Hypoxia Intervention

An altitude simulator unit (Sierra 100, Hypoxico Altitude Training Systems, Gardiner, NY, United States) was used to modulate FiO_2_ and deliver the hypoxic stimulus (Figure 1A). Participants were required to wear a naso-oral facemask for the entirety of both visits. Following baseline measures at sea level, the ACH intervention was delivered over a 15-minute titration phase to control the rate of exposure (.01 FiO_2_ reduction every 1.5 minutes). During the AIH intervention, participants did not experience the 15-minute titration phase, instead they were exposed immediately to the hypoxic stimulus. The cyclic nature of the AIH intervention was achieved by utilising two Hans Rudolph three-way stopcock values to exhaust the hypoxia stimulus back into the atmosphere and expose the participant to the ambient room air. The SpO_2_ of each participant was monitored using a finger pulse oximeter (Berry, Shanghai Berry Electronic Tech Co., Ltd., Minhang District, Shanghai, China) and was the primary measure to quantify the individual’s experience of hypoxia. As the sensitivity of SpO_2_ is varied across and within populations (Rojas-Camayo et al., 2018), SpO_2_ was not clamped throughout the intervention and allowed to move dynamically in response to the hypoxia interventions. Participants remained in the hypoxic stimulus for 1 hr in both the ACH and AIH interventions.

**Figure 1.**
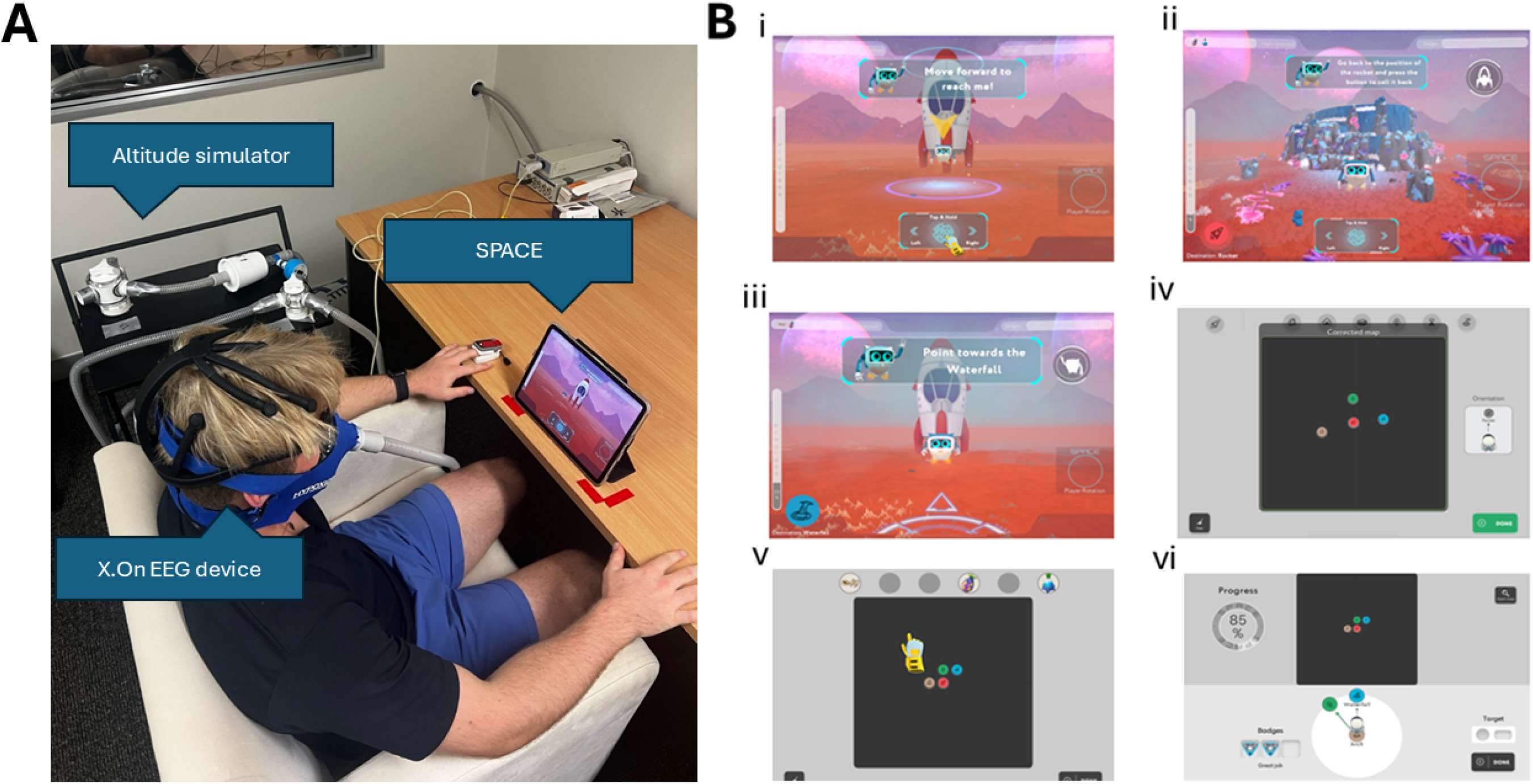
Participant set up and each phase of the Spatial Assessment for Cognitive Evaluation (SPACE). Participants sat comfortably in an upright position for the duration of both visits (A). A full facemask was attached to the participant before baseline measurements were taken and was used to deliver the hypoxic and sham intervention. Peripheral blood oxygen saturation (SpO_2_) was continuously monitored using a finger pulse oximeter throughout the 1 hour of exposure. Participants performed the navigational task on an iPad positioned 50 cm in front of them (B). SPACE consists of 6 phases that are selected purposefully for distinguishing between good and bad navigators. These consist of a training (Bi), path integration (Bii), egocentric pointing (Biii), mapping (Biv), associative memory (Bv), and perspective taking (Bvi). Pictured is the author Dylan Stenholm.

### Navigational task

Participants performed a battery of navigational tasks from SPACE that were delivered on an iPad. SPACE is a serious VR game that is readily used as a digital assessment tool to characterise individual differences in navigational ability and cognitive impairment in both healthy and clinical populations (Colombo et al., 2024; Colombo, Minta, Taylor, et al., 2026; Colombo, Minta, Thrash, et al., 2026; Colombo, 2023; McKeown et al., 2026; Tian et al., 2026). Each task is specifically designed to assess key components of navigational ability that are used in everyday navigation. These five tasks consist of path integration, egocentric pointing, mapping, associative memory, and perspective-taking.

#### Training phase

Participants first completed a training phase to familiarise themselves with the navigation controls (e.g., translation and rotation). Guided by an animated robot, L15A, participants practised these movements by following simple routes through the virtual environment before progressing to the assessment tasks (Figure 1Bi).

#### Path integration

Participants begin at the rocket, located in the centre of the environment, and follow the robot to two landmarks where items are collected. After reaching the second landmark, participants are asked to navigate back to the rocket without guidance, relying on their ability to continuously update changes in position and orientation (Figure 1Bii).

#### Egocentric pointing

Following the path integration task, participants complete a series of pointing trials. They are first asked to indicate the direction of each previously visited landmark from the rocket. Participants are then positioned at one of the landmarks and required to point towards the remaining landmark and the rocket (Figure 1Biii).

#### Mapping

Participants are presented with a top-down view of the environment and asked to reconstruct the spatial layout by placing the landmarks in their correct locations relative to one another and to the rocket (Figure 1Biv).

#### Associative memory

Participants are asked to recall which item was collected at each landmark during the path integration task. Participants are presented with a map of the environment and asked to drag and drop each item onto the landmark where it was originally encountered (Figure 1Bv).

#### Perspective taking

Participants complete a series of perspective-taking trials using a top-down view of the environment. For each trial, they are instructed to imagine standing at one landmark while facing a second landmark and then indicate the direction of a third landmark relative to that imagined viewpoint (Figure 1Bvi).

### EEG Acquisition and Preprocessing

EEG was recorded before and following the sham or hypoxia sessions for ACH and AIH interventions. Five minutes of resting-state eyes-closed (EC) and 5 minutes of eyes-open (EO) EEG were recorded from a subset of the participants (ACH = 18, AIH = 22). Seven channels of EEG were recorded using a portable EEG system (X.on EEG Headset) using sponge-based electrodes bathed in a saline solution. Electrodes were organised according to the 10-20 EEG system positioned at F3, F4, C3, Cz, C4, P3, and P4. An additional electrode was attached to the right earlobe for grounding and online referencing. EEG was sampled at 250Hz with a 50 Hz notch filter. EEG data were exported as BrainVision files for preprocessing.

Preprocessing of the EEG data was conducted in Python (v3.11.5) using MNE package (v1.5.1). EC and EO EEG recordings were processed individually. Signals were high-passed filtered at .1Hz to remove any low-frequency drift in the signal and then re-referenced to the average of all electrodes. Independent Component Analysis (ICA) was applied to identify artifacts, using four components, the “infomax” method, and the maximum number of iterations set to “auto”. MNE ICALabel was used to classify components as either “brain”, “muscle”, “heartbeat”, or “other”. Only “brain” and “other” components were retained for reconstruction. Following ICA corrections, the EEG signal was segmented into 2000 ms epochs. AutoReject was used to identify artefact-contaminated epochs, such as those containing blink artifact, and were removed. Following this, epochs that had greater than ± 150 µV deviation were also removed. Power spectral densities were calculated for each single 2000ms segment and then averaged into a single power spectral density using a Welch’s method, with 300 points used for Fast Fourier Transformation with 50% overlap. The spectra were calculated over the frequency range from 1 Hz to the Nyquist frequency (125 Hz) to avoid aliasing.

The *SpecParam* algorithm was used for separation of periodic (parameterised power and central frequency) and aperiodic (exponent and offset of the broadband) EEG activity from the power spectral densities. The *SpecParam* models were fit between 3 – 40 Hz range and consisted of the following parameters: peak width limit of 1 – 8 Hz, a minimum peak height of 0.01, a peak threshold of 2.0, maximum of infinite peaks, and the aperiodic mode set to “fixed”. The aperiodic exponent, offset, and model fit (R^2^) were obtained from the model outputs. The mean fit of the models across all participants, sessions, and electrode locations was 0.97 ± 0.06.

Alpha and beta central frequency and power were quantified following parameterisation using *SpecParam*. The exponent and offset of the power spectral densities were quantified to reflect aperiodic EEG activity. Participants that had less than 50% of channels with missing data were replaced with the mean of the other channels for the relative session and timepoint. If participants had more than 50% of channels with missing data, they were excluded from the analysis for that relative EEG variable. No participants had greater than 50% of channels missing for alpha of beta activity, nor the aperiodic exponent and offset. Each EEG variable was then averaged across all seven electrodes to create a scalp-wide composite measure.

### Survey assessments

Participants completed two surveys that characterise their perceived navigational ability and their experience of adverse symptoms to the hypoxic stimulus.

#### The Santa Barbara Sense of Direction Scale (SBSOD)

The SBSOD is a self-reported questionnaire consisting of 15-items presented as Likert scales pertaining to the individual’s perceived spatial ability (Hegarty et al., 2002). Participants are required to select their response on a scale between 1 “Strongly agree” to 7 “Strongly disagree” to statements such as “I am very good at judging distances” and “I don’t have a very good ‘mental map’ of my environment”. Final scores were computed by averaging item responses, with higher scores indicating a stronger self-perceived sense of direction. The SBSOD scale has demonstrated high internal consistencies (α = .97 - .88; Davies et al., 2017; Zafeiriou et al., 2021) and excellent test-retest reliability (r = .91; Hegarty et al., 2002). There were no significant differences in self-perceived navigational ability between our intervention groups (ACH *Md* = 4.13 ± 1.23, AIH *Md* = 4.40 ± .93; *U* = 247*, p* = .298).

#### The Lake Louise Acute Mountain Sickness Score (LLAMSS)

The LLAMSS is a self-reported survey consisting of four rated symptoms that characterise acute mountain sickness (AMS) of which participants must rate their perceived experience (Roach et al., 2018). These consist of headache, nausea/gastrointestinal symptoms, fatigue and/or weakness, and dizziness/light-headedness. Participants rate these symptoms as 0 “None at all”, 1 “Mild”, 2 “Moderate”, or 3 “Severe”. Scores for each symptom are then summed, and an accumulative score is recorded. Accumulative scores are categorized as mild AMS (3 – 5 points), moderate AMS (6 – 9 points), and severe (10 – 12 points). The scale has demonstrated high internal consistency (α = .84; Macinnis et al., 2013). There were no significant differences in LLAMS between our intervention groups (ACH *Md* = 0.00 ± 3.00, AIH *Md* = 0.00 ± 1.00; *U* = 383.50*, p* = .127).

### Experimental procedures

Participants attended the laboratory on two separate occasions where they performed the hypoxia or sham session on their initial visit. All procedures were the same between interventions except that .209 FiO_2_ was maintained throughout the sham session but was reduced to .13 FiO_2_. In the ACH intervention, the target FiO_2_ was reached by progressively reducing .015 FiO_2_ over a 15 min titration phase. Conversely during the AIH intervention, the target FiO_2_ exposure was presented immediately at the onset of the intermittent cycles.

SBSODs and baseline measures of SpO_2,_ heart rate, EEG, and SPACE were performed before the titration phase. Immediately following the titration phase, SpO_2_ and heart rate measures were recorded again, and then 30 minutes and 1 hour following the titration phase. Once participants were exposed to 1 hour of the intervention, EEG and SPACE performance were recorded again.

### Data analysis and reduction

#### SBSOD

Items 1,3,4,5,7,9, and 14 were first reversed scored and then all 15 items were summed then an average score between 1 and 7 was calculated with 1 meaning a poor sense of direction and 7 meaning a good sense of direction.

#### SpO_2_ and Heart Rate

For each timepoint measurement of SpO_2_ and heart rate during the ACH intervention, recordings were averaged over a 2-minute period to represent SpO_2_ for the timepoint. For each cycle measurement of SpO_2_ and heart rate during the AIH intervention, seven cycle blocks were created by averaging the 2-minute cycles in groups of five. These consisted of Block 0 (Premeasures), Block 1 (0 – 10 minutes), Block 2 (10 – 20 minutes), Block 3 (20 – 30 minutes), Block 4 (30 – 40 minutes), Block 5 (40 – 50 minutes), and Block 6 (50 – 60 minutes).

#### SPACE

Seven trials of the Path Integration task were performed where completion time (sec) and distance error in game units (1 unit = 50 cm) were averaged across all trials. Nine trials of the Egocentric Pointing task were performed where absolute pointing angle error was averaged across trials. Participants only performed one iteration of the Mapping task where the *R^2^* of the estimated and corrected map were calculated using bidimensional regression (Friedman & Kohler, 2003; Tobler, 1994). Similarly, one iteration of the Associative Memory task was performed where percentage of correct associations were calculated. Seven trials of the Perspective-Taking task were performed where angle error were averaged across all trials.

### Statistical analysis

All statistical analyses were performed in R (version 2025.05.1). Topographic EEG maps were generated using MNE-Python. For the SPACE and EEG measures, values were normalised to baseline to account for inter-day variability in navigation performance and brain activity. Change scores were calculated as the difference between post-intervention and baseline values. Due to a ceiling effect in the Associative Memory task (100% accuracy across all participants and sessions), this measure was excluded from further analyses.

Non-parametric factorial analyses were conducted using the aligned rank transform (ART) procedure implemented in the ARTool package (Kay et al., 2025; Wobbrock et al., 2011). The ART procedure permits analysis of factorial designs using standard ANOVA while accommodating violations of normality. Separate mixed-effects ART ANOVAs were fitted for each outcome measure, with participant included as a random intercept to account for repeated measures.

To examine physiological responses to hypoxia, ART ANOVAs included the fixed effects of Session (Sham, Hypoxia), Time (ACH) or Block (AIH), and their interaction. To test Hypothesis 1 (acute hypoxia impairs navigation), ART ANOVAs examined the effects of Session (Sham, Hypoxia), Intervention (ACH, AIH), and their interaction on SPACE performance. To test Hypothesis 2 (acute hypoxia alters resting-state EEG), ART ANOVAs examined the effects of Session (Sham, Hypoxia), Intervention (ACH, AIH), Eye State (eyes closed, eyes open), and their interactions on periodic and aperiodic EEG measures. To test Hypothesis 3 (inter-individual differences in oxygen desaturation predict behavioural and electrophysiological responses), linear regression models were fitted using only the hypoxia sessions, with SpO□, Intervention (ACH, AIH), and their interaction included as predictors.

Significant main effects and interactions were followed by post hoc pairwise comparisons using estimated marginal means with Tukey-adjusted *p* values. Statistical significance was set at *p* < .05.

## RESULTS

### Physiological response to hypoxia

For SpO_2_ during the ACH protocol, there was a main effect of Session, *F*(1,336) = 1194.61, *p* < .001, where SpO_2_ was lower during the hypoxia session compared to the sham session. There was a main effect of Timepoint, *F*(3,336) = 188.92, *p* < .001. Pairwise contrasts revealed that SpO_2_ was lower at the 0 min, 30 min, and 60 min (all *p* <.001) timepoints compared to Premeasures. There was a Session by Time interaction, *F*(3,336) = 170.45, *p* < .001. Pairwise contrasts revealed that SpO_2_ during the hypoxia session was lower at 0 min, 30 min, and 60 min (all *p* <.001) compared to Premeasures but not during the sham session (Figure 2A). For heart rate during the ACH protocol, there was a main effect of Session, *F*(1,336) = 5.93, *p* < .015, where heart rate was higher during the hypoxia session compared to the sham session. There was a main effect of Timepoint, *F*(1,336) = 8.56, *p* < .001. Pairwise contrasts revealed that by 60 mins heart rate was higher than Premeasures (*p* < .001).

**Figure 2.**
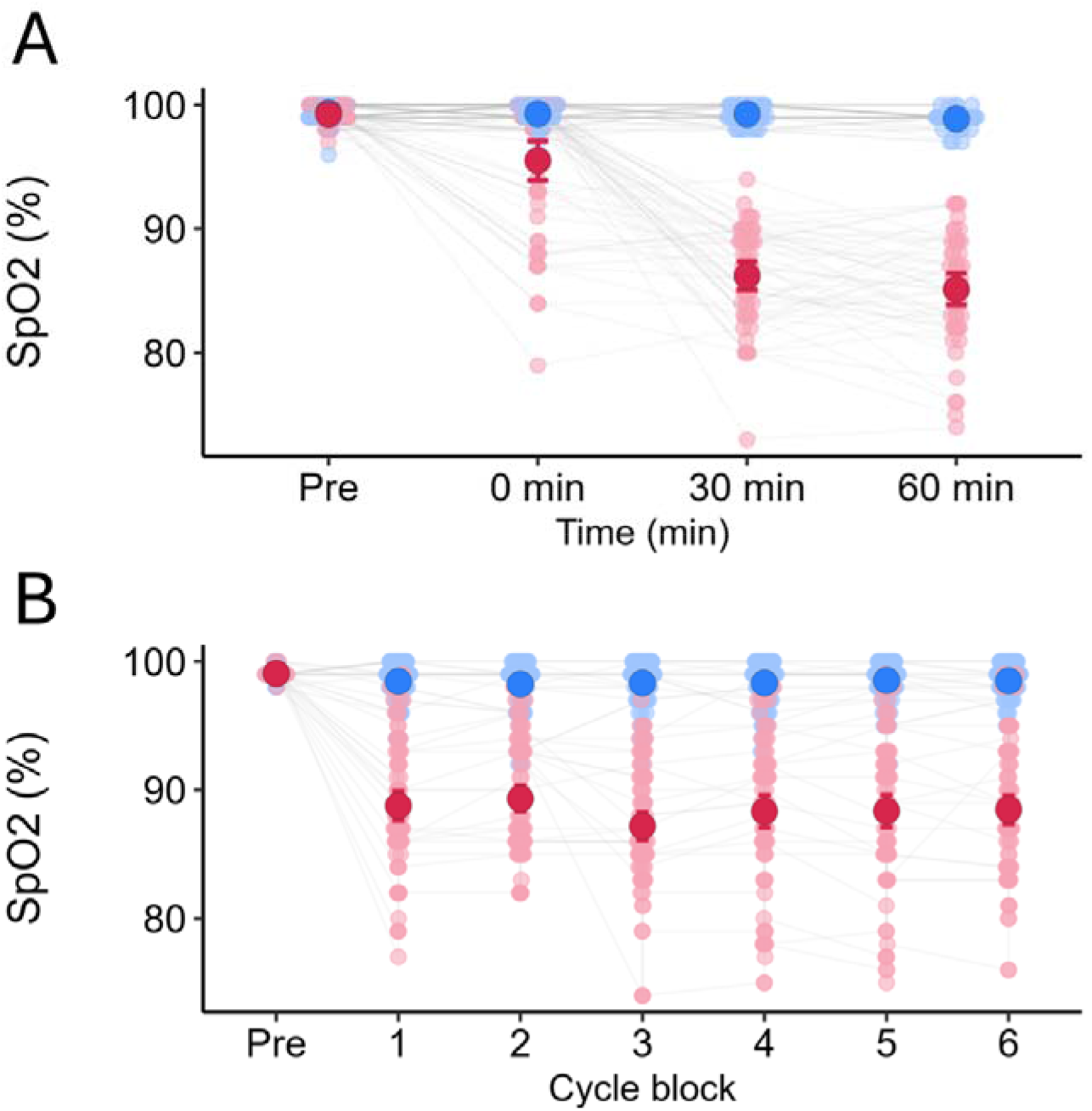
Physiological responses during the acute continuous hypoxia (ACH; A) and acute intermittent hypoxia (AIH: B) interventions. During ACH, SpO□ was lower during the hypoxia session (red circles) compared with sham (blue circles) and remained suppressed across all timepoints (0, 0.5, and 1 h), reflected by significant main effects of session and timepoint and a significant session by timepoint interaction. During AIH, SpO□ was lower during the hypoxia session compared with sham across hypoxic cycle blocks, reflected by significant main effects of session and block and a significant session by block interaction. Figures show estimated marginal means and error bars represent 95% confidence intervals. ACH, acute continuous hypoxia; AIH, acute intermittent hypoxia; SpO□, peripheral oxygen saturation.

For SpO_2_ during the AIH protocol, there was a main effect of Session, *F*(1,963) = 1608.58, *p* < .001, where SpO_2_ was lower during the hypoxia session compared to the sham session. There was a main effect of Block, *F*(6,963) = 17.05, *p* < .001. Pairwise contrasts revealed that SpO_2_ was lower at Block 1 to 6 (all *p* < .001) compared to Premeasures. There was a Session by Block interaction, *F*(6,963) = 16.29, *p* < .001. Pairwise contrasts revealed that SpO_2_ during the hypoxia but not the sham session was lower at Block 1 to 6 (all *p* < .001) compared to Premeasures (Figure 2B). For heart rate during the ACH protocol, there was a main effect of Session, *F*(1,897.23) = 17.09, *p* = .015, where heart rate was higher during the hypoxia session compared to the sham session. There was a main effect of Block, *F*(6,897.11) = 5.56, *p* < .001. Pairwise contrasts revealed that by Block 6 heart rate was higher than Premeasures (*p* < .001).

### Navigation performance

#### Path integration

For time to complete the path integration task, there was no main effect of Intervention, *F*(1,71.26) = .66, *p* = .419, Session, *F*(1,210.07) = 2.35, *p* = .127, nor an Intervention by Session interaction, *F*(1,209.73) = .45, *p* = .505. For distance error, there was a main effect of Session, *F*(1,215.76) = 14.17, *p* < .001, where distance error was greater during the hypoxia session compared to the sham session (Figure 3A). There was no main effect of Intervention, *F*(1,71.63) = .65, *p* = .423, nor a Session by Intervention interaction, *F*(1,215.40) = .55, *p* =.459. There was no association between the reduction in SpO□ during hypoxia and change in path integration distance error (Supplementary Material).

**Figure 3.**
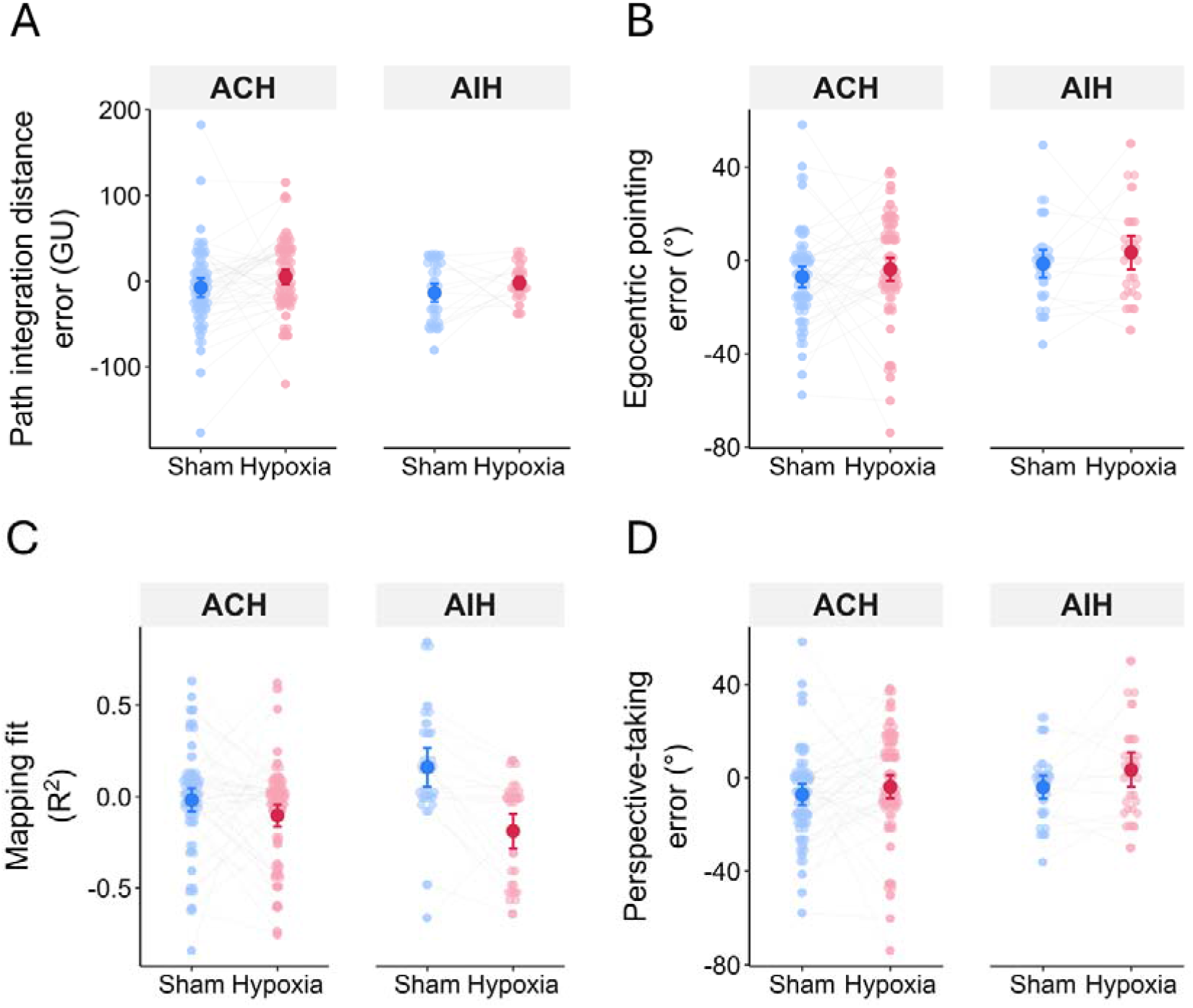
Change from baseline in Spatial Performance Assessment for Cognitive Evaluation (SPACE) performance during sham (blue) and hypoxia (red) sessions following acute continuous (ACH) and intermittent hypoxia (AIH). Session main effects occurred for all outcomes, with hypoxia increasing path integration distance error (A) and egocentric pointing error (B). Session by intervention interactions occurred for mapping fit (C) and perspective taking error (D), with poorer mapping during hypoxia in AIH and greater perspective taking error in ACH. Symbols show estimated marginal means with 95% confidence intervals.

#### Egocentric pointing

For angle error during the egocentric pointing task, there was a main effect of Session, *F*(1,210.96) = 5.87, *p* = .016, where error was greater during the hypoxia session compared to the sham session (Figure 3B). There was no main effect of Intervention, *F*(1,71.10) = 1.02, *p* = .316, nor a Session by Intervention interaction, *F*(1,210.97) = .06, *p* = .811. The association between absolute SpO□ and change in egocentric pointing error differed between ACH and AIH groups (SpO□ × Intervention: β = −2.13, SE = 1.06, *t*(128) = −2.01, *p* = .046). However, simple slopes were not significant within either ACH (β = 0.49, 95% CI [−0.57, 1.54]) or AIH (β = −1.64, 95% CI [−3.45, 0.17]) groups.

#### Mapping

For correctness of mapping fit, there was a main effect of Session, *F*(1,208.75) = 14.04, *p* < .001, where fit was worse during the hypoxia session compared to the sham session. There was no main effect of Intervention, *F*(1,70.92) = 2.98, *p* = .089, but there was a Session by Intervention interaction, *F*(1,208.80) = 10.65, *p* < .001, where hypoxia worsened mapping fit more in the AIH intervention group (*p* < .001) compared to the ACH intervention group (*p* = .051; Figure 3C). There was no association between the reduction in SpO□ during hypoxia and change in mapping fit (Supplementary Material).

#### Perspective taking

For error during the perspective taking task, there was a main effect of Session, *F*(1,214.03) = 11.80, *p* < .001, where perspective taking angle error was greater during hypoxia compared to sham. There was no main effect of Intervention, *F*(1,71.72) = .04, *p* = .844, but there was a Session by Intervention interaction, *F*(1,214.50) = 20.27, *p* < .001, where hypoxia had a greater impact on perspective taking error in the ACH group (*p* < .001) compared to the AIH group (*p* = .053). There was no association between the reduction in SpO□ during hypoxia and change in perspective taking error (Supplementary Material).

### Resting-state EEG

#### Aperiodic EEG activity

For the aperiodic EEG exponent, there was a main effect of Session, *F*(1,106.24) = 5.05, *p* = .027, where the exponent was reduced in the hypoxia session compared to the sham session (Figure 4A). There was no main effect of Intervention, *F*(1,36.91) < 001, *p* = .977, but a Session by Intervention interaction, *F*(1,105.78) = 8.85, *p* = .004, where the exponent was reduced more in hypoxia for the AIH group (*p* = .001) but did not reduce in the ACH group (*p* = .981). There was no main effect of Eye State, *F*(1,89.77) = .01, *p* = .927, nor an Eye State interaction with Session, *F*(1,88.86) = 1.74, *p* = .191, Intervention, *F*(1,89.93) = 3.02, *p* = .085, nor their three-way interaction, *F*(1,88.84) = .47, *p* = .496. There were no significant effects for the aperiodic offset (see Supplementary Material). There were no associations between the reduction in SpO□ during hypoxia and change in the aperiodic exponent nor the offset (Supplementary Material).

**Figure 4.**
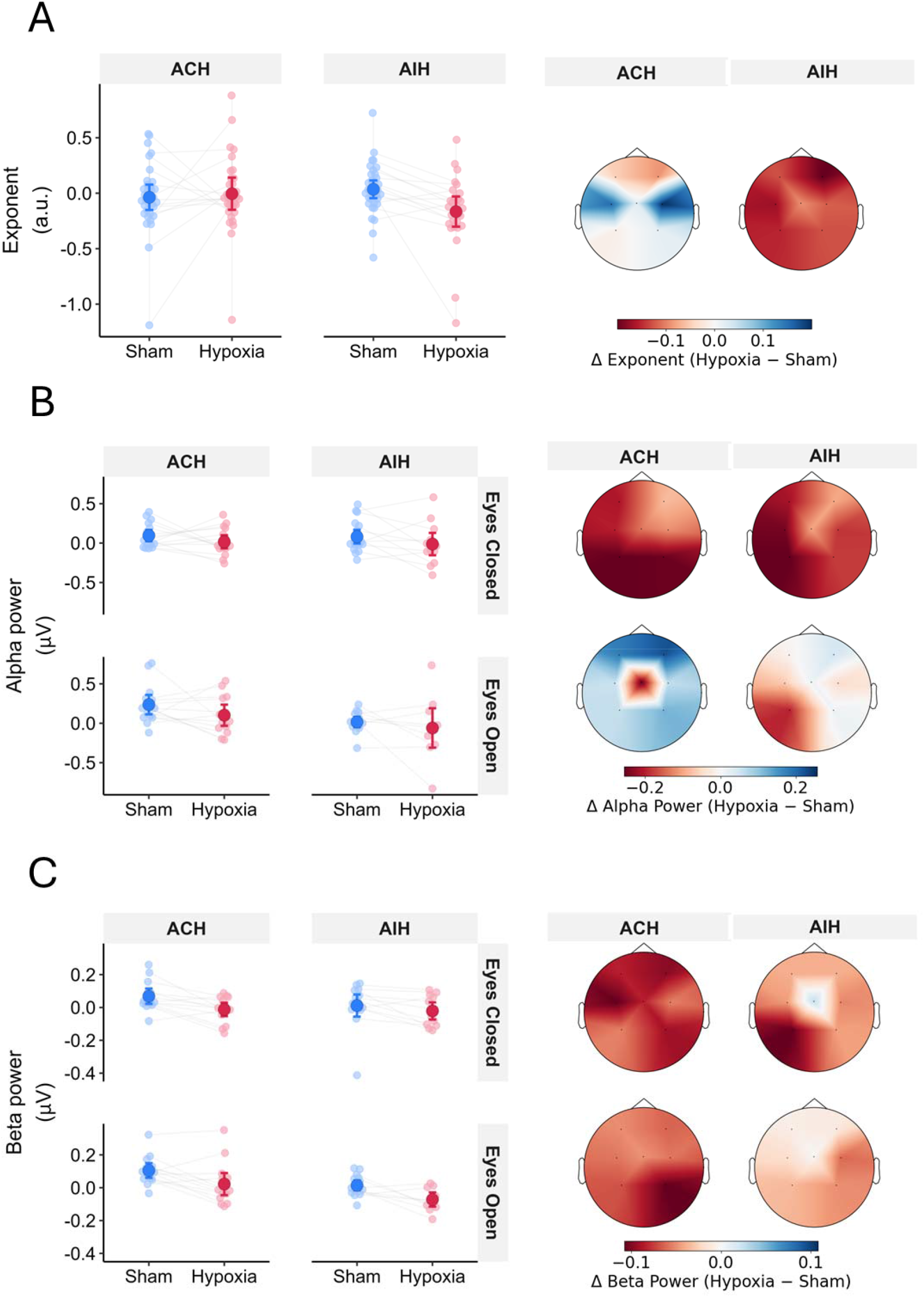
Aperiodic exponent (A), alpha power (B), and beta power (C) during sham (blue) and hypoxia (red) sessions following acute continuous (ACH) and intermittent hypoxia (AIH). Hypoxia reduced aperiodic exponent, with a greater reduction following AIH but not ACH. Alpha and beta power decreased during hypoxia without session by intervention interactions. Intervention by eye state interactions reflected lower eyes-open alpha and beta power in AIH than ACH. Connected points represent individual participants across sessions. Larger symbols show estimated marginal means with 95% confidence intervals. Scalp maps show hypoxia minus sham differences.

#### Periodic EEG activity

For alpha power, there was a main effect of Session, *F*(1,91.33) = 10.84, *p* = .001, where alpha power was smaller during the hypoxia session compared to the sham session. There was a main effect of Intervention, *F*(1,36.53) = 4.53, *p* = .040, where alpha power was lower in the AIH group compared to the ACH group. There was no Session by Intervention interaction, *F*(1,90.98) = .22, *p* = .644. There was also no main effect of Eye State, *F*(1,81.88) = 1.73, *p* = .192, nor interaction with Session, *F*(1,78.21) = .26, *p* = .609, but there was an interaction effect with Intervention, *F*(1,82.24) = 4.74, *p* < .032, where eyes-open alpha power was lower in the AIH group compared to the ACH group (*p* = .034; Figure 4B). There was no three-way interaction, *F*(1,78.12) = .06, *p* = .810. There were no effects for alpha central frequency (see Supplementary Material). There were no associations between the reduction in SpO□ during hypoxia and change in the alpha power nor central frequency (Supplementary Material).

For beta power, there was a main effect of Session, *F*(1,83.28) = 25.68, *p* < .001, where beta power was lower in the hypoxia session compared to the sham session (Figure 4C). There was no main effect of Intervention, *F*(1,33.45) = 3.82, *p* = .059, nor a Session by Intervention interaction, *F*(1,82.52) = .24, *p* = .624. There was no main effect of Eye State, *F*(1,73.36) < .01, *p* = .947, interaction with Session, *F*(1,72.29) = .06, *p* = .416, but there was an interaction with Intervention, *F*(1,73.27) = 4.05, *p* = .048, where EO but not EC beta power was lower in the AIH group compared to the ACH group (*p* = .034; Figure 4C). For beta central frequency, there was a main effect of Session, *F*(1,89.60) = 4.39, *p* = .048. There were no other effects for beta central frequency (see Supplementary Material). There were no associations between the reduction in SpO□ during hypoxia and change in the beta power nor central frequency (Supplementary Material).

## DISCUSSION

The current study investigated how acute hypoxic exposure affects navigational performance and EEG-derived measures of intrinsic brain activity. Participants completed SPACE and resting-state EEG recordings before and after 1 h of ACH or AIH. Acute hypoxia impaired performance across all SPACE tasks, including path integration, egocentric pointing, mapping, and perspective taking, although the magnitude of impairment for some outcomes differed between ACH and AIH. Hypoxia also altered multiple features of the resting EEG, including reductions in the aperiodic exponent and in alpha- and beta-oscillatory power. The reduction in the aperiodic exponent was particularly evident for the AIH group, whereas reductions in alpha and beta oscillatory power were observed across both interventions. Together, these findings suggest that acute hypoxia can disrupt multiple components of navigation while altering EEG-derived measures of intrinsic brain activity. We propose that these alterations in EEG characteristics may reflect changes in the capacity of neural systems to coordinate information processing under hypoxic stress, which coincides with impairments in navigation.

### Physiological responses to hypoxia

Both the ACH and AIH interventions produced substantial reductions in SpO_2_, indicating that our experimental manipulations effectively induced hypoxia. During ACH, SpO_2_ decreased steadily over the 1 hr exposure. Conversely, AIH produced cyclic reductions in SpO_2_ that followed a profile of transient hypoxic exposure interspersed with normoxic air. Heart rate remained largely stable during ACH but showed a progressive increase across cycles during AIH. This likely reflects autonomic adjustments to alternating normoxic air and hypoxic air states (Iturriaga et al., 2014). While both interventions produced similar overall reductions in SpO_2_, the temporal differences allowed us to examine whether the pattern of exposure differentially affects intrinsic brain activity and navigation.

Despite temporal differences, both interventions produced similar, broad impairments in navigation ability and alterations in intrinsic brain activity, specifically in path integration and egocentric pointing, as well as in alpha and beta power. However, the pattern of hypoxic delivery appeared to influence specific outcomes. Mapping performance was reduced to a greater extent during AIH compared to ACH. Similarly, perspective taking was worse during ACH compared to AIH. The aperiodic exponent was also reduced during AIH but not during ACH. This delivery-specific pattern also contrasts with our original hypothesis that AIH would improve navigation performance. This hypothesis was based on evidence that continuous exposure to hypoxia impairs cognitive function, whereas repeated brief intervals of hypoxic exposure may induce hormetic and neuroplastic adaptations, particularly in some clinical and older populations (Boulares et al., 2024; Burtscher et al., 2024; Schega et al., 2013, 2016). Instead, AIH produced impairments across several navigational outcomes and to a greater extent in some instances, with mapping performance being mostly affected. These findings suggest that a single 1-hour exposure may be insufficient to elicit the adaptive effects reported following repeated AIH interventions. Thus, while the absolute severity of hypoxic exposure may be sufficient to produce general impairments across both interventions, the temporal pattern of oxygen availability may shape the specific cognitive and neural processes that are most affected.

### Acute hypoxia disrupts multiple processes supporting spatial navigation

Acute hypoxia impaired multiple components of navigation, as reflected by increased errors in path integration, egocentric pointing, mapping, and perspective taking. However, the pattern of impairment did not remain consistent across all tasks or interventions. Path integration distance error and egocentric pointing angle error increased during hypoxia, regardless of whether hypoxia was delivered continuously or intermittently. This suggests that the ability to spatially update is sensitive to acute reductions in oxygen availability, irrespective of the temporal pattern. Alternatively, mapping performance and perspective taking demonstrated temporally specific effects of hypoxia. Here, the ability to reconstruct a map of the environment was particularly impaired during AIH, while the ability to shift between flexible representations of the environment was particularly impaired during ACH. Together, these findings indicate that acute hypoxia does not impair navigational processes to the same degree, as some tasks showed impairment that depended on the temporal pattern of exposure.

The impairment observed across both allocentric, and egocentric components of SPACE challenges a simple account in which acute hypoxia selectively disrupts hippocampal-dependent, place-based navigation. Instead, the results suggest that acute hypoxia may affect multiple interacting systems that support spatial behaviour, consistent with the distributed organisation of navigation (Baumann & Mattingley, 2021; Byrne et al., 2007; Chrastil et al., 2015; Dhindsa et al., 2014; Ekstrom et al., 2005, 2017; R. A. Epstein et al., 2017; Howard et al., 2014; O’Keefe, 1991; Sahoo et al., 2026; Sherrill et al., 2015; Wolbers & Buchel, 2005). Rather than indicating vulnerability confined to a single navigational process, our findings suggest that acute hypoxia can compromise multiple processes involved in constructing, maintaining, updating, retrieving, and transforming spatial representations. This interpretation is consistent with neurophysiological evidence that multiple distributed brain networks, particularly hippocampal and entorhinal circuits, are vulnerable to hypoxia (Biswal et al., 2016; Hencz et al., 2023; Lu et al., 2025; Micaux et al., 2025; Shushanyan et al., 2023; Vetrovoy et al., 2021; Zhang et al., 2018; Zhu et al., 2019).

Larger errors in path integration demonstrates this oxygen dependence clearly and is consistent with our original hypothesis that navigational processes relying heavily on hippocampal-entorhinal circuitry may be particularly vulnerable to reduced oxygen availability. Successful path integration requires continuous updating of position from visual and self-motion cues which place sustained demands on systems involved in spatial representation (Angelaki & Cullen, 2008; Ekstrom & Ranganath, 2018; Etienne & Jeffery, 2004). The increase in distance error observed under both interventions suggests that this process is sensitive to acute reductions in oxygen availability, regardless of whether hypoxia is sustained or intermittent. Previous work indicates that hippocampal function is particularly sensitive to hypoxic stress, capable of altering CA3-CA1 synaptic dynamics, NMDA receptor activity and neuromodulatory signalling (Ai & Baker, 2006; Bekker et al., 2006; Biswal et al., 2016; Gibson & Duffy, 1981; Ji et al., 2021; Kumral et al., 2004; Maiti et al., 2008; Quintana et al., 2015). Such alterations could compromise the neural computations required to maintain and update spatial representations over time. Indeed, this impairment may have been amplified in the present study, since path integration was performed without physical movement, thereby limiting vestibular and proprioceptive input and potentially increasing reliance on visual processes that are initially impaired by hypoxic stress (Blacker et al., 2021; Blacker & McHail, 2022; Connolly et al., 2008).

The effects on mapping and perspective taking tasks further suggest that hypoxia disrupts the manipulation of spatial representations but is dependent on the temporal profile of exposure (Baumann & Mattingley, 2021; Byrne et al., 2007; Dhindsa et al., 2014; Ekstrom et al., 2017; Epstein et al., 2017; Sahoo et al., 2026; Sherrill et al., 2015; Wolbers & Buchel, 2005). Mapping performance deteriorated to a greater extent during AIH compared to ACH, while perspective taking performance deteriorated to a greater extent during ACH compared to AIH. Mapping requires the construction and maintenance of a configurational representation of environmental landmarks and boundaries, whereas perspective taking requires the transformation of spatial representation from different viewpoints. Both tasks depend on distributed neural networks that extend beyond the hippocampus, including parahippocampal, retrosplenial, and striatal regions (Baumann & Mattingley, 2021; Ekstrom & Hill, 2023; Iaria et al., 2003). The divergent effects of ACH and AIH may therefore be explained by differences in the temporal demand of hypoxic stress imposed on these supporting networks by the ACH and AIH interventions. Here, continuous hypoxia may preferentially challenge sustained spatial transformation, whereas intermittent hypoxia may disrupt the repeated construction and updating of abstract spatial representations held in short-term memory. Although this interpretation remains mechanistic, it is consistent with evidence that the effects of hypoxia on broader cognition depend on both the severity and temporal characteristics of exposure (Asmaro et al., 2013; Bondi et al., 2021; McMorris et al., 2017; Turner, Barker-Collo, et al., 2015). Future research should aim to elucidate this phenomenon further.

The increase in egocentric pointing error extends this interpretation beyond navigation processes traditionally considered to depend strongly on hippocampal-entorhinal networks.

Egocentric navigation can recruit broader cortical and striatal systems, including parahippocampal, retrosplenial and caudate regions (Baumann & Mattingley, 2021; Ekstrom & Hill, 2023; Iaria et al., 2003). The presence of impairment in egocentric pointing alongside impairment in allocentric navigation suggests that the effects of acute hypoxia are unlikely to be restricted to a single navigational strategy. The egocentric pointing task requires participants to draw on allocentrically derived spatial information and orient themselves with reference to landmarks from an egocentric perspective. This suggests that hypoxia has a global effect of reducing the capacity of neural networks to support the continuum of navigational strategies required for navigation (Ekstrom et al., 2017). The absence of a Session by Intervention interaction suggests that egocentric pointing was similarly affected across the two exposure profiles, indicating that acute hypoxic stress was sufficient to impair this component of navigation irrespective of whether hypoxia was continuous or intermittent.

Together, our behavioural findings suggest that acute hypoxia compromises many processes of navigation, rather than selectively impairing one specific strategy. The differential effects observed in our behavioural findings following ACH and AIH further indicate that the temporal dynamics of oxygen availability may influence which navigational strategies are most vulnerable to hypoxic stress. Furthermore, hypoxia likely compromised the efficiency of neural networks supporting navigation and may explain the differentiation in behavioural effects between the two interventions. This possibility provides a framework for interpreting the accompanying changes observed in intrinsic EEG activity.

### Hypoxia-related alterations in intrinsic brain activity

The impairment observed in navigation was accompanied by selective alterations in EEG measures of intrinsic brain activity. Acute hypoxia reduced alpha and beta power and was associated with a flatter aperiodic exponent, while the aperiodic offset remained unchanged. Hypoxia also reduced beta central frequency, although this effect was comparatively modest. Together, these findings suggest that acute reductions in oxygen availability alter multiple features of ongoing cortical activity.

Reductions in both alpha and beta power provide converging evidence that hypoxia alters oscillatory mechanisms implicated in global neural coordination (Chang et al., 2022; Feddersen et al., 2007; Hutcheon et al., 2023; Otto et al., 2025; Rice et al., 2019a, 2019b; Zhao et al., 2016). Alpha oscillations reflect the regulation of information processing, attentional control, and sensory gating, whereas beta activity has been linked to the maintenance and coordination of distributed cognitive representations (Engel & Fries, 2010; Klimesch, 2012; MacLean et al., 2012; Schmidt et al., 2019). This interpretation may be particularly relevant to navigation, which depends on coordinated activity of distributed hippocampal, entorhinal, retrosplenial, parahippocampal and cortical systems. Successful path integration, egocentric updating, map construction and flexibility in perspective require information to be integrated across these partially overlapping networks. Consequently, a hypoxia-related reduction in the efficiency or stability of large-scale neural network coordination may affect several navigational processes concurrently, consistent with the broad pattern of behavioural impairment observed in the present study. However, this explanation remains inferential because the EEG was recorded at rest rather than during active navigation. Future research combining hypoxic exposure with navigation-based EEG assessments is required to determine if changes in navigation-specific oscillatory indices of neural dynamics are evident.

The reduction in the aperiodic exponent provides complementary evidence that hypoxia also alters underlying spectral features of neural networks. The exponent reflects the slope of the aperiodic component of the power spectrum and has been associated with features of population-level neural dynamics and excitation–inhibition balance, although its precise physiological interpretation remains debated (Donoghue et al., 2020; Gao et al., 2017; Preston et al., 2026). In the current study, the exponent was reduced overall during hypoxia, with the effect driven primarily by a larger reduction following AIH. This pattern suggests that acute hypoxia also alters the broader balance of neural dynamics across frequencies. Although reductions in the exponent have been proposed to reflect changes in excitation–inhibition balance, the present findings cannot directly establish the degree to which changes in excitation or inhibition contribute to the global hypoxic-disruption to neural dynamics (Donoghue et al., 2020; Gao et al., 2017; Preston et al., 2026). Nonetheless, the concurrent reductions in exponent, alpha power, and beta power suggest that hypoxia affects multiple components of the resting EEG spectrum, consistent with a broader alteration in intrinsic neural dynamics rather than an isolated change in a single oscillatory mechanism.

The stronger reduction in exponent during AIH further indicates that the temporal pattern of hypoxia may shape neurophysiological responses. The differences in the exposure profiles of ACH and AIH may impose distinct demands on the cerebrovascular, autonomic, and neural systems (Ainslie & Subudhi, 2014; Burtscher et al., 2024; Iturriaga et al., 2014). Our observation of an intervention-specific difference in the exponent is consistent with the possibility that fluctuating oxygen availability processes may produce a distinct pattern of neurophysiological stress. Notably, this finding did not support our original expectation that AIH would increase aperiodic activity in a manner consistent with enhanced neural plasticity. Although repeated intermittent hypoxic exposure has been proposed to promote adaptive and neuroplastic responses in some instances (Boulares et al., 2024; Burtscher et al., 2024; Schega et al., 2013, 2016) but see Champod et. al. (2013), the present acute AIH protocol instead produced a larger reduction in the aperiodic exponent. This distinction may reflect differences between the acute neurophysiological effects of a single intermittent hypoxic exposure and the longer-term adaptations that emerge following repeated AIH protocols. As no interaction effects were observed for oscillatory measures, the temporal profile of hypoxia may modulate aspects of intrinsic neural activity without fundamentally altering the broader reduction of oscillatory power.

Interestingly, hypoxia had relatively limited effects on oscillatory frequency and the aperiodic offset. Although beta central frequency was reduced during hypoxia, this effect was comparatively modest relative to the reductions observed in alpha and beta power, while alpha central frequency and aperiodic offset remained unchanged. This pattern suggests that the hypoxic exposure used in the present study did not produce a uniform slowing of cortical oscillations or a generalised shift in broadband spectral activity. Instead, acute hypoxia appeared to preferentially affect the amplitude and spectral organisation of ongoing neural activity. This differs from the more pronounced EEG slowing occasionally reported during severe or prolonged hypoxic exposures (Chang et al., 2022; Feddersen et al., 2007; Zhao et al., 2016). One possibility is that the moderate and relatively brief exposure used here was sufficient to alter oscillatory amplitude and broader spectral dynamics without producing more extensive changes in oscillatory frequency (Nakata et al., 2025; Otto et al., 2025).

Together, the EEG findings suggest that acute hypoxia alters intrinsic brain activity across both periodic and aperiodic components of the EEG spectrum. The concurrent reduction in alpha and beta power and the aperiodic exponent is consistent with a broader disruption to the organisation of ongoing neural activity, potentially affecting the capacity of neural systems to coordinate information processing. This systems-level interpretation provides a plausible explanation for the widespread impairment that was observed across the SPACE tasks, although a direct mechanistic relationship between our resting state EEG and navigational deficits cannot be established.

Notably, the magnitude of peripheral oxygen desaturation was not associated with changes in either EEG measures or navigational performance in the present study. Although previous research has identified associations between oxygen saturation and cognitive impairment under some hypoxic conditions, these relationships appear to vary according to exposure severity and the cognitive outcome assessed (Hohenauer et al., 2022; McMorris et al., 2017). Similarly, electrophysiological responses to hypoxia appear to depend on the severity, duration, and pattern of oxygen delivery, rather than reflecting uniform changes in spectral power or generalised slowing. This variability suggests that the effects of hypoxia on brain activity may involve changes in specific neural processes supporting information processing across distributed brain systems, rather than being determined solely by the degree of peripheral oxygen desaturation. Accordingly, behavioural and neurophysiological responses may also depend on cerebral oxygen delivery, cerebrovascular regulation, autonomic responses, and other compensatory mechanisms (Ainslie & Poulin, 2004; Ainslie & Subudhi, 2014; Hoiland et al., 2018). Future studies combining peripheral oxygen saturation with direct measures of cerebral oxygenation and cerebrovascular function may help clarify the physiological factors underlying individual differences in hypoxia tolerance.

### Limitations

The hypoxic exposures used in the present study were moderate in severity and relatively short in duration. Consequently, the magnitude and pattern of behavioural and neurophysiological effects observed here may not reflect the full extent of impairment that could occur during more severe or prolonged hypoxic exposures, including those encountered in operational, environmental, or clinical contexts (Coronel-Oliveros et al., 2023; Crişan et al., 2014). Future studies should therefore examine whether the effects observed in the present study scale with the severity and duration of hypoxic exposure.

The use of a 7-channel EEG system limited the spatial resolution of the present study and may have reduced sensitivity to more localised or region-specific effects of hypoxia. This limitation is particularly relevant to the interpretation of aperiodic activity, as changes in the aperiodic exponent to hypoxia may vary across cortical regions (Coronel-Oliveros et al., 2023). Future studies using higher-density EEG may provide greater insight into the spatial distribution of hypoxia-related changes in periodic and aperiodic activity. In addition, resting-state EEG was measured separately from the SPACE tasks, limiting the extent to which the observed changes in intrinsic brain activity can be directly linked to individual differences in navigational performance. Recording EEG during navigation tasks could provide greater insight into the neural processes associated with the maintenance or disruption of spatial performance under hypoxic conditions.

A further limitation concerns the sex distribution of the sample, which comprised 63 females and nine males. Large-scale research and meta-analytic evidence indicate an average male advantage in some spatial navigation tasks, although the magnitude of this difference varies across populations, task demands, and outcome measures (García-Navarra et al., 2026; Nazareth et al., 2019; Spiers et al., 2023). The predominantly female sample may therefore limit the generalisability of our findings across sex groups. Moreover, the small number of male participants precluded a robust examination of whether sex moderated the behavioural or neurophysiological effects of hypoxia. Future studies with more balanced samples are needed to establish whether sex influences susceptibility to hypoxia-related changes in spatial navigation and intrinsic brain activity.

Participants may have relied on different cognitive strategies to complete the SPACE tasks. Differences in strategy use, perceived task difficulty, and the relatively limited motor demands of the tasks may have contributed to variability in performance across participants. Importantly, the SPACE tasks were completed without physical movement through the environment, limiting vestibular and proprioceptive contributions to navigation and increasing reliance on visual and internally generated spatial information (Baumann & Mattingley, 2021). Although alterations in visuoperceptual processing and visual attention may contribute to cognitive impairment during hypoxic exposure (Blacker et al., 2021; Blacker & McHail, 2022), the absence of physical movement limits the generalisability of the present findings to real-world navigation. Future studies incorporating immersive or physically active navigation paradigms could therefore determine whether the effects observed here extend to more ecologically valid forms of spatial behaviour.

Importantly, this finding did not support our third hypothesis that individuals with larger reductions in SpO□ would demonstrate greater impairment in navigation and intrinsic brain activity. The absence of these associations suggests that peripheral oxygen desaturation alone may not adequately capture individual vulnerability to hypoxia. Instead, the functional consequences of hypoxia may depend on interactions among oxygen delivery, cerebrovascular compensation, autonomic responses, neural resilience, cerebral blood flow, carbon dioxide, and ventilation (Ainslie & Subudhi, 2014). The absence of these measures, therefore, limits our ability to determine whether differences between ACH and AIH reflected direct neural effects of the temporal exposure profile, cerebrovascular compensation, autonomic adaptation, or broader systemic responses. Future studies incorporating multimodal physiological monitoring will be important for identifying the mechanisms underlying individual and exposure-specific responses to acute hypoxia.

## Conclusion

This study demonstrates that acute hypoxia impairs multiple components of navigational performance while producing selective alterations in intrinsic brain activity. Performance was impaired across all SPACE tasks, with greater path integration and egocentric pointing errors observed during hypoxia irrespective of intervention type, whereas the effects on mapping and perspective taking differed according to the temporal pattern of hypoxic exposure. These findings indicate that acute hypoxia can disrupt both allocentric and egocentric aspects of navigation while suggesting that continuous and intermittent hypoxia may differentially affect specific navigation processes. Hypoxia also altered intrinsic EEG activity, with reductions in aperiodic exponent, alpha power, beta power, and beta central frequency, whereas aperiodic offset and alpha central frequency remained largely stable. The reduction in aperiodic exponent was particularly evident following AIH, while intervention-related differences in EO alpha and beta power further suggest that the temporal pattern of oxygen delivery may influence specific aspects of intrinsic neural activity. Importantly, the magnitude of peripheral oxygen desaturation was not associated with changes in navigational performance or EEG activity, suggesting that SpO□ alone may not fully capture individual vulnerability to the cognitive and neurophysiological effects of hypoxia. Together, these findings demonstrate that even relatively brief, moderate hypoxic exposure can impair navigation and alter intrinsic neural dynamics, with the temporal pattern of oxygen availability moderating some, but not all, behavioural and neurophysiological responses.

## CODE AVAILABILITY

Data and scripts for the analysis of the current study are available at https://github.com/MindSpaceLab/OXYNAV.

