## Supplementary Material 1 for "Acute Hypoxia Disrupts Multiple Processes Supporting Spatial Navigation": Supplementary Material 1.html (from another version of this preprint: version 1, 2026-09-30)

ART ANOVA OXYNAV


### ART ANOVA OXYNAV

#### 2026-08-03

CONTINUOUS SPO2

```
## Warning: package 'ARTool' was built under R version 4.5.3
```

```
## NOTE: Results may be misleading due to involvement in interactions
```

```
##  contrast            estimate   SE  df t.ratio p.value
##  Session0 - Session1      194 5.61 336  34.563  <.0001
## 
## Results are averaged over the levels of: Timepoint 
## Degrees-of-freedom method: kenward-roger
```

```
## NOTE: Results may be misleading due to involvement in interactions
```

```
##  contrast                estimate   SE  df t.ratio p.value
##  Timepoint1 - Timepoint2     60.3 10.3 336   5.846  <.0001
##  Timepoint1 - Timepoint3    185.1 10.3 336  17.956  <.0001
##  Timepoint1 - Timepoint4    209.8 10.3 336  20.351  <.0001
##  Timepoint2 - Timepoint3    124.8 10.3 336  12.109  <.0001
##  Timepoint2 - Timepoint4    149.5 10.3 336  14.505  <.0001
##  Timepoint3 - Timepoint4     24.7 10.3 336   2.396  0.0799
## 
## Results are averaged over the levels of: Session 
## Degrees-of-freedom method: kenward-roger 
## P value adjustment: tukey method for comparing a family of 4 estimates
```

```
##  contrast  estimate   SE  df t.ratio p.value
##  0,1 - 0,2     9.84 13.4 336   0.735  0.9959
##  0,1 - 0,3    14.35 13.4 336   1.071  0.9623
##  0,1 - 0,4    53.70 13.4 336   4.010  0.0019
##  0,1 - 1,1    15.48 13.4 336   1.156  0.9435
##  0,1 - 1,2    72.73 13.4 336   5.431  <.0001
##  0,1 - 1,3   212.74 13.4 336  15.887  <.0001
##  0,1 - 1,4   220.42 13.4 336  16.460  <.0001
##  0,2 - 0,3     4.51 13.4 336   0.337  1.0000
##  0,2 - 0,4    43.87 13.4 336   3.276  0.0254
##  0,2 - 1,1     5.64 13.4 336   0.421  0.9999
##  0,2 - 1,2    62.90 13.4 336   4.697  0.0001
##  0,2 - 1,3   202.91 13.4 336  15.152  <.0001
##  0,2 - 1,4   210.58 13.4 336  15.725  <.0001
##  0,3 - 0,4    39.36 13.4 336   2.939  0.0682
##  0,3 - 1,1     1.13 13.4 336   0.085  1.0000
##  0,3 - 1,2    58.39 13.4 336   4.360  0.0005
##  0,3 - 1,3   198.40 13.4 336  14.815  <.0001
##  0,3 - 1,4   206.07 13.4 336  15.388  <.0001
##  0,4 - 1,1   -38.22 13.4 336  -2.854  0.0855
##  0,4 - 1,2    19.03 13.4 336   1.421  0.8471
##  0,4 - 1,3   159.04 13.4 336  11.876  <.0001
##  0,4 - 1,4   166.71 13.4 336  12.449  <.0001
##  1,1 - 1,2    57.26 13.4 336   4.275  0.0007
##  1,1 - 1,3   197.27 13.4 336  14.731  <.0001
##  1,1 - 1,4   204.94 13.4 336  15.304  <.0001
##  1,2 - 1,3   140.01 13.4 336  10.455  <.0001
##  1,2 - 1,4   147.68 13.4 336  11.028  <.0001
##  1,3 - 1,4     7.67 13.4 336   0.573  0.9992
## 
## Degrees-of-freedom method: kenward-roger 
## P value adjustment: tukey method for comparing a family of 8 estimates
```

```
## Warning: package 'ggbeeswarm' was built under R version 4.5.3
```

```
## 
## Attaching package: 'dplyr'
```

```
## The following objects are masked from 'package:stats':
## 
##     filter, lag
```

```
## The following objects are masked from 'package:base':
## 
##     intersect, setdiff, setequal, union
```

```
## Warning: package 'ggnewscale' was built under R version 4.5.3
```

CONTINUOUS HR

```
## NOTE: Results may be misleading due to involvement in interactions
```

```
##  contrast            estimate   SE  df t.ratio p.value
##  Session0 - Session1    -19.4 7.96 336  -2.434  0.0154
## 
## Results are averaged over the levels of: Timepoint 
## Degrees-of-freedom method: kenward-roger
```

```
## NOTE: Results may be misleading due to involvement in interactions
```

```
##  contrast                estimate   SE  df t.ratio p.value
##  Timepoint1 - Timepoint2      2.9 11.1 336   0.261  0.9938
##  Timepoint1 - Timepoint3    -26.5 11.1 336  -2.383  0.0823
##  Timepoint1 - Timepoint4    -45.5 11.1 336  -4.093  0.0003
##  Timepoint2 - Timepoint3    -29.4 11.1 336  -2.644  0.0425
##  Timepoint2 - Timepoint4    -48.4 11.1 336  -4.354  0.0001
##  Timepoint3 - Timepoint4    -19.0 11.1 336  -1.710  0.3201
## 
## Results are averaged over the levels of: Session 
## Degrees-of-freedom method: kenward-roger 
## P value adjustment: tukey method for comparing a family of 4 estimates
```

INTERMITTENT SPO2

```
## NOTE: Results may be misleading due to involvement in interactions
```

```
##  contrast            estimate   SE  df t.ratio p.value
##  Session0 - Session1      464 11.6 963  40.107  <.0001
## 
## Results are averaged over the levels of: Block 
## Degrees-of-freedom method: kenward-roger
```

```
## NOTE: Results may be misleading due to involvement in interactions
```

```
##  contrast        estimate   SE  df t.ratio p.value
##  Block0 - Block1  408.512 49.2 963   8.308  <.0001
##  Block0 - Block2  393.131 49.2 963   7.995  <.0001
##  Block0 - Block3  492.669 49.2 963  10.020  <.0001
##  Block0 - Block4  413.622 49.2 963   8.412  <.0001
##  Block0 - Block5  426.212 49.2 963   8.668  <.0001
##  Block0 - Block6  425.872 49.2 963   8.661  <.0001
##  Block1 - Block2  -15.381 28.4 963  -0.542  0.9982
##  Block1 - Block3   84.156 28.4 963   2.964  0.0486
##  Block1 - Block4    5.109 28.4 963   0.180  1.0000
##  Block1 - Block5   17.700 28.4 963   0.623  0.9961
##  Block1 - Block6   17.359 28.4 963   0.612  0.9965
##  Block2 - Block3   99.537 28.4 963   3.506  0.0086
##  Block2 - Block4   20.491 28.4 963   0.722  0.9913
##  Block2 - Block5   33.081 28.4 963   1.165  0.9069
##  Block2 - Block6   32.741 28.4 963   1.153  0.9111
##  Block3 - Block4  -79.047 28.4 963  -2.784  0.0798
##  Block3 - Block5  -66.456 28.4 963  -2.341  0.2256
##  Block3 - Block6  -66.797 28.4 963  -2.353  0.2201
##  Block4 - Block5   12.591 28.4 963   0.444  0.9994
##  Block4 - Block6   12.250 28.4 963   0.432  0.9995
##  Block5 - Block6   -0.341 28.4 963  -0.012  1.0000
## 
## Results are averaged over the levels of: Session 
## Degrees-of-freedom method: kenward-roger 
## P value adjustment: tukey method for comparing a family of 7 estimates
```

```
##  contrast  estimate   SE  df t.ratio p.value
##  0,0 - 0,1   45.806 40.8 963   1.124  0.9977
##  0,0 - 0,2   59.000 40.8 963   1.447  0.9760
##  0,0 - 0,3   50.100 40.8 963   1.229  0.9945
##  0,0 - 0,4   57.050 40.8 963   1.399  0.9820
##  0,0 - 0,5   46.256 40.8 963   1.135  0.9975
##  0,0 - 0,6   45.144 40.8 963   1.107  0.9980
##  0,0 - 1,0    1.219 52.6 963   0.023  1.0000
##  0,0 - 1,1  509.050 40.8 963  12.486  <.0001
##  0,0 - 1,2  501.619 40.8 963  12.304  <.0001
##  0,0 - 1,3  562.956 40.8 963  13.808  <.0001
##  0,0 - 1,4  514.312 40.8 963  12.615  <.0001
##  0,0 - 1,5  515.606 40.8 963  12.647  <.0001
##  0,0 - 1,6  517.194 40.8 963  12.686  <.0001
##  0,1 - 0,2   13.194 23.5 963   0.561  1.0000
##  0,1 - 0,3    4.294 23.5 963   0.182  1.0000
##  0,1 - 0,4   11.244 23.5 963   0.478  1.0000
##  0,1 - 0,5    0.450 23.5 963   0.019  1.0000
##  0,1 - 0,6   -0.662 23.5 963  -0.028  1.0000
##  0,1 - 1,0  -44.587 40.8 963  -1.094  0.9983
##  0,1 - 1,1  463.244 23.5 963  19.680  <.0001
##  0,1 - 1,2  455.812 23.5 963  19.364  <.0001
##  0,1 - 1,3  517.150 23.5 963  21.970  <.0001
##  0,1 - 1,4  468.506 23.5 963  19.904  <.0001
##  0,1 - 1,5  469.800 23.5 963  19.959  <.0001
##  0,1 - 1,6  471.387 23.5 963  20.026  <.0001
##  0,2 - 0,3   -8.900 23.5 963  -0.378  1.0000
##  0,2 - 0,4   -1.950 23.5 963  -0.083  1.0000
##  0,2 - 0,5  -12.744 23.5 963  -0.541  1.0000
##  0,2 - 0,6  -13.856 23.5 963  -0.589  1.0000
##  0,2 - 1,0  -57.781 40.8 963  -1.417  0.9799
##  0,2 - 1,1  450.050 23.5 963  19.120  <.0001
##  0,2 - 1,2  442.619 23.5 963  18.804  <.0001
##  0,2 - 1,3  503.956 23.5 963  21.410  <.0001
##  0,2 - 1,4  455.312 23.5 963  19.343  <.0001
##  0,2 - 1,5  456.606 23.5 963  19.398  <.0001
##  0,2 - 1,6  458.194 23.5 963  19.466  <.0001
##  0,3 - 0,4    6.950 23.5 963   0.295  1.0000
##  0,3 - 0,5   -3.844 23.5 963  -0.163  1.0000
##  0,3 - 0,6   -4.956 23.5 963  -0.211  1.0000
##  0,3 - 1,0  -48.881 40.8 963  -1.199  0.9957
##  0,3 - 1,1  458.950 23.5 963  19.498  <.0001
##  0,3 - 1,2  451.519 23.5 963  19.182  <.0001
##  0,3 - 1,3  512.856 23.5 963  21.788  <.0001
##  0,3 - 1,4  464.212 23.5 963  19.721  <.0001
##  0,3 - 1,5  465.506 23.5 963  19.776  <.0001
##  0,3 - 1,6  467.094 23.5 963  19.844  <.0001
##  0,4 - 0,5  -10.794 23.5 963  -0.459  1.0000
##  0,4 - 0,6  -11.906 23.5 963  -0.506  1.0000
##  0,4 - 1,0  -55.831 40.8 963  -1.369  0.9851
##  0,4 - 1,1  452.000 23.5 963  19.203  <.0001
##  0,4 - 1,2  444.569 23.5 963  18.887  <.0001
##  0,4 - 1,3  505.906 23.5 963  21.493  <.0001
##  0,4 - 1,4  457.262 23.5 963  19.426  <.0001
##  0,4 - 1,5  458.556 23.5 963  19.481  <.0001
##  0,4 - 1,6  460.144 23.5 963  19.548  <.0001
##  0,5 - 0,6   -1.113 23.5 963  -0.047  1.0000
##  0,5 - 1,0  -45.038 40.8 963  -1.105  0.9981
##  0,5 - 1,1  462.794 23.5 963  19.661  <.0001
##  0,5 - 1,2  455.363 23.5 963  19.345  <.0001
##  0,5 - 1,3  516.700 23.5 963  21.951  <.0001
##  0,5 - 1,4  468.056 23.5 963  19.885  <.0001
##  0,5 - 1,5  469.350 23.5 963  19.940  <.0001
##  0,5 - 1,6  470.938 23.5 963  20.007  <.0001
##  0,6 - 1,0  -43.925 40.8 963  -1.077  0.9985
##  0,6 - 1,1  463.906 23.5 963  19.708  <.0001
##  0,6 - 1,2  456.475 23.5 963  19.393  <.0001
##  0,6 - 1,3  517.812 23.5 963  21.998  <.0001
##  0,6 - 1,4  469.169 23.5 963  19.932  <.0001
##  0,6 - 1,5  470.462 23.5 963  19.987  <.0001
##  0,6 - 1,6  472.050 23.5 963  20.054  <.0001
##  1,0 - 1,1  507.831 40.8 963  12.456  <.0001
##  1,0 - 1,2  500.400 40.8 963  12.274  <.0001
##  1,0 - 1,3  561.737 40.8 963  13.778  <.0001
##  1,0 - 1,4  513.094 40.8 963  12.585  <.0001
##  1,0 - 1,5  514.388 40.8 963  12.617  <.0001
##  1,0 - 1,6  515.975 40.8 963  12.656  <.0001
##  1,1 - 1,2   -7.431 23.5 963  -0.316  1.0000
##  1,1 - 1,3   53.906 23.5 963   2.290  0.5627
##  1,1 - 1,4    5.263 23.5 963   0.224  1.0000
##  1,1 - 1,5    6.556 23.5 963   0.279  1.0000
##  1,1 - 1,6    8.144 23.5 963   0.346  1.0000
##  1,2 - 1,3   61.337 23.5 963   2.606  0.3379
##  1,2 - 1,4   12.694 23.5 963   0.539  1.0000
##  1,2 - 1,5   13.988 23.5 963   0.594  1.0000
##  1,2 - 1,6   15.575 23.5 963   0.662  1.0000
##  1,3 - 1,4  -48.644 23.5 963  -2.067  0.7242
##  1,3 - 1,5  -47.350 23.5 963  -2.012  0.7603
##  1,3 - 1,6  -45.763 23.5 963  -1.944  0.8015
##  1,4 - 1,5    1.294 23.5 963   0.055  1.0000
##  1,4 - 1,6    2.881 23.5 963   0.122  1.0000
##  1,5 - 1,6    1.587 23.5 963   0.067  1.0000
## 
## Degrees-of-freedom method: kenward-roger 
## P value adjustment: tukey method for comparing a family of 14 estimates
```

INTERMITTENT HR

```
## NOTE: Results may be misleading due to involvement in interactions
```

```
##  contrast            estimate   SE  df t.ratio p.value
##  Session0 - Session1    -68.2 16.5 897  -4.134  <.0001
## 
## Results are averaged over the levels of: Block 
## Degrees-of-freedom method: kenward-roger
```

```
## NOTE: Results may be misleading due to involvement in interactions
```

```
##  contrast        estimate   SE  df t.ratio p.value
##  Block0 - Block1  -146.38 41.5 897  -3.524  0.0081
##  Block0 - Block2  -171.78 41.7 897  -4.116  0.0008
##  Block0 - Block3  -144.29 41.7 897  -3.463  0.0100
##  Block0 - Block4   -90.71 41.7 897  -2.176  0.3095
##  Block0 - Block5   -98.77 41.6 897  -2.372  0.2117
##  Block0 - Block6  -174.96 41.8 897  -4.185  0.0006
##  Block1 - Block2   -25.41 24.7 897  -1.028  0.9475
##  Block1 - Block3     2.08 24.6 897   0.085  1.0000
##  Block1 - Block4    55.67 24.6 897   2.258  0.2657
##  Block1 - Block5    47.60 24.6 897   1.937  0.4562
##  Block1 - Block6   -28.59 24.9 897  -1.150  0.9123
##  Block2 - Block3    27.49 25.0 897   1.102  0.9276
##  Block2 - Block4    81.07 25.0 897   3.247  0.0206
##  Block2 - Block5    73.01 24.9 897   2.931  0.0535
##  Block2 - Block6    -3.18 25.2 897  -0.126  1.0000
##  Block3 - Block4    53.58 24.9 897   2.156  0.3210
##  Block3 - Block5    45.52 24.8 897   1.836  0.5240
##  Block3 - Block6   -30.67 25.1 897  -1.223  0.8851
##  Block4 - Block5    -8.07 24.8 897  -0.325  0.9999
##  Block4 - Block6   -84.26 25.1 897  -3.362  0.0141
##  Block5 - Block6   -76.19 25.0 897  -3.047  0.0383
## 
## Results are averaged over the levels of: Session 
## Degrees-of-freedom method: kenward-roger 
## P value adjustment: tukey method for comparing a family of 7 estimates
```

```
##  contrast   estimate   SE  df t.ratio p.value
##  0,0 - 0,1  -79.9125 58.1 897  -1.376  0.9845
##  0,0 - 0,2 -141.6062 58.1 897  -2.438  0.4534
##  0,0 - 0,3 -100.8500 58.1 897  -1.736  0.9026
##  0,0 - 0,4  -80.7500 58.1 897  -1.390  0.9830
##  0,0 - 0,5  -79.8688 58.1 897  -1.375  0.9845
##  0,0 - 0,6 -129.4688 58.1 897  -2.229  0.6084
##  0,0 - 1,0   -9.6562 75.0 897  -0.129  1.0000
##  0,0 - 1,1 -211.5133 58.5 897  -3.616  0.0218
##  0,0 - 1,2 -204.3304 59.0 897  -3.461  0.0367
##  0,0 - 1,3 -188.6709 58.9 897  -3.205  0.0803
##  0,0 - 1,4 -105.6876 58.9 897  -1.794  0.8787
##  0,0 - 1,5 -116.8602 58.8 897  -1.988  0.7753
##  0,0 - 1,6 -217.5814 59.3 897  -3.672  0.0180
##  0,1 - 0,2  -61.6938 33.5 897  -1.840  0.8575
##  0,1 - 0,3  -20.9375 33.5 897  -0.624  1.0000
##  0,1 - 0,4   -0.8375 33.5 897  -0.025  1.0000
##  0,1 - 0,5    0.0437 33.5 897   0.001  1.0000
##  0,1 - 0,6  -49.5562 33.5 897  -1.478  0.9714
##  0,1 - 1,0   70.2562 58.1 897   1.209  0.9953
##  0,1 - 1,1 -131.6008 34.2 897  -3.845  0.0096
##  0,1 - 1,2 -124.4179 35.2 897  -3.538  0.0284
##  0,1 - 1,3 -108.7584 34.9 897  -3.118  0.1024
##  0,1 - 1,4  -25.7751 34.9 897  -0.738  1.0000
##  0,1 - 1,5  -36.9477 34.7 897  -1.063  0.9987
##  0,1 - 1,6 -137.6689 35.5 897  -3.875  0.0086
##  0,2 - 0,3   40.7563 33.5 897   1.215  0.9951
##  0,2 - 0,4   60.8563 33.5 897   1.815  0.8694
##  0,2 - 0,5   61.7375 33.5 897   1.841  0.8569
##  0,2 - 0,6   12.1375 33.5 897   0.362  1.0000
##  0,2 - 1,0  131.9500 58.1 897   2.272  0.5766
##  0,2 - 1,1  -69.9070 34.2 897  -2.042  0.7404
##  0,2 - 1,2  -62.7242 35.2 897  -1.784  0.8832
##  0,2 - 1,3  -47.0647 34.9 897  -1.349  0.9869
##  0,2 - 1,4   35.9187 34.9 897   1.028  0.9991
##  0,2 - 1,5   24.7461 34.7 897   0.712  1.0000
##  0,2 - 1,6  -75.9752 35.5 897  -2.139  0.6742
##  0,3 - 0,4   20.1000 33.5 897   0.599  1.0000
##  0,3 - 0,5   20.9812 33.5 897   0.626  1.0000
##  0,3 - 0,6  -28.6187 33.5 897  -0.853  0.9999
##  0,3 - 1,0   91.1937 58.1 897   1.570  0.9534
##  0,3 - 1,1 -110.6633 34.2 897  -3.233  0.0740
##  0,3 - 1,2 -103.4804 35.2 897  -2.943  0.1618
##  0,3 - 1,3  -87.8209 34.9 897  -2.518  0.3967
##  0,3 - 1,4   -4.8376 34.9 897  -0.138  1.0000
##  0,3 - 1,5  -16.0102 34.7 897  -0.461  1.0000
##  0,3 - 1,6 -116.7314 35.5 897  -3.286  0.0633
##  0,4 - 0,5    0.8812 33.5 897   0.026  1.0000
##  0,4 - 0,6  -48.7188 33.5 897  -1.453  0.9752
##  0,4 - 1,0   71.0938 58.1 897   1.224  0.9947
##  0,4 - 1,1 -130.7633 34.2 897  -3.820  0.0105
##  0,4 - 1,2 -123.5804 35.2 897  -3.514  0.0308
##  0,4 - 1,3 -107.9209 34.9 897  -3.094  0.1093
##  0,4 - 1,4  -24.9376 34.9 897  -0.714  1.0000
##  0,4 - 1,5  -36.1102 34.7 897  -1.039  0.9990
##  0,4 - 1,6 -136.8314 35.5 897  -3.851  0.0094
##  0,5 - 0,6  -49.6000 33.5 897  -1.479  0.9712
##  0,5 - 1,0   70.2125 58.1 897   1.209  0.9953
##  0,5 - 1,1 -131.6446 34.2 897  -3.846  0.0096
##  0,5 - 1,2 -124.4617 35.2 897  -3.539  0.0283
##  0,5 - 1,3 -108.8022 34.9 897  -3.119  0.1020
##  0,5 - 1,4  -25.8188 34.9 897  -0.739  1.0000
##  0,5 - 1,5  -36.9914 34.7 897  -1.065  0.9987
##  0,5 - 1,6 -137.7127 35.5 897  -3.876  0.0085
##  0,6 - 1,0  119.8125 58.1 897   2.063  0.7269
##  0,6 - 1,1  -82.0446 34.2 897  -2.397  0.4833
##  0,6 - 1,2  -74.8617 35.2 897  -2.129  0.6812
##  0,6 - 1,3  -59.2022 34.9 897  -1.697  0.9168
##  0,6 - 1,4   23.7812 34.9 897   0.681  1.0000
##  0,6 - 1,5   12.6086 34.7 897   0.363  1.0000
##  0,6 - 1,6  -88.1127 35.5 897  -2.480  0.4231
##  1,0 - 1,1 -201.8570 58.5 897  -3.451  0.0378
##  1,0 - 1,2 -194.6742 59.0 897  -3.297  0.0611
##  1,0 - 1,3 -179.0147 58.9 897  -3.041  0.1260
##  1,0 - 1,4  -96.0313 58.9 897  -1.630  0.9380
##  1,0 - 1,5 -107.2039 58.8 897  -1.823  0.8653
##  1,0 - 1,6 -207.9252 59.3 897  -3.509  0.0313
##  1,1 - 1,2    7.1829 35.8 897   0.201  1.0000
##  1,1 - 1,3   22.8424 35.5 897   0.643  1.0000
##  1,1 - 1,4  105.8257 35.6 897   2.971  0.1508
##  1,1 - 1,5   94.6531 35.4 897   2.673  0.2965
##  1,1 - 1,6   -6.0681 36.2 897  -0.168  1.0000
##  1,2 - 1,3   15.6595 36.4 897   0.430  1.0000
##  1,2 - 1,4   98.6428 36.5 897   2.703  0.2787
##  1,2 - 1,5   87.4702 36.3 897   2.408  0.4749
##  1,2 - 1,6  -13.2510 37.1 897  -0.357  1.0000
##  1,3 - 1,4   82.9833 36.2 897   2.294  0.5601
##  1,3 - 1,5   71.8107 36.0 897   1.994  0.7715
##  1,3 - 1,6  -28.9105 36.8 897  -0.786  1.0000
##  1,4 - 1,5  -11.1726 36.0 897  -0.310  1.0000
##  1,4 - 1,6 -111.8939 36.7 897  -3.046  0.1244
##  1,5 - 1,6 -100.7212 36.6 897  -2.753  0.2510
## 
## Degrees-of-freedom method: kenward-roger 
## P value adjustment: tukey method for comparing a family of 14 estimates
```

PATH INTEGRATION TIME TO COMPLETE

```
## 
## Call:
## lm(formula = average_pi_total_time_change ~ spo2 * intervention, 
##     data = hx_dat)
## 
## Residuals:
##      Min       1Q   Median       3Q      Max 
## -31.5577  -0.8817   2.3810   4.9518  12.2179 
## 
## Coefficients:
##                               Estimate Std. Error t value Pr(>|t|)
## (Intercept)                    27.4174    17.2886   1.586    0.115
## spo2                           -0.3137     0.2027  -1.547    0.124
## interventionintermittent      -52.4541    34.8734  -1.504    0.135
## spo2:interventionintermittent   0.5747     0.4008   1.434    0.154
## 
## Residual standard error: 8.996 on 132 degrees of freedom
## Multiple R-squared:  0.04152,    Adjusted R-squared:  0.01974 
## F-statistic: 1.906 on 3 and 132 DF,  p-value: 0.1317
```

PATH INTEGRATION DISTANCE ERROR

```
## NOTE: Results may be misleading due to involvement in interactions
```

```
##  contrast          estimate   SE  df t.ratio p.value
##  (ses-0) - (ses-1)    -31.2 8.28 216  -3.764  0.0002
## 
## Results are averaged over the levels of: intervention 
## Degrees-of-freedom method: kenward-roger
```

```
## 
## Call:
## lm(formula = average_pi_distance_change ~ spo2 * intervention, 
##     data = hx_dat)
## 
## Residuals:
##      Min       1Q   Median       3Q      Max 
## -125.770  -23.161   -2.535   27.375  109.696 
## 
## Coefficients:
##                               Estimate Std. Error t value Pr(>|t|)
## (Intercept)                   -17.5205    76.8700  -0.228    0.820
## spo2                            0.2647     0.9013   0.294    0.769
## interventionintermittent      -33.4454   155.0574  -0.216    0.830
## spo2:interventionintermittent   0.2976     1.7822   0.167    0.868
## 
## Residual standard error: 40 on 132 degrees of freedom
## Multiple R-squared:  0.007414,   Adjusted R-squared:  -0.01514 
## F-statistic: 0.3287 on 3 and 132 DF,  p-value: 0.8046
```

EGOCENTRIC POINTING ERROR

```
## Welcome to emmeans.
## Caution: You lose important information if you filter this package's results.
## See '? untidy'
```

```
## NOTE: Results may be misleading due to involvement in interactions
```

```
##  contrast          estimate   SE  df t.ratio p.value
##  (ses-0) - (ses-1)    -18.9 7.81 211  -2.424  0.0162
## 
## Results are averaged over the levels of: intervention 
## Degrees-of-freedom method: kenward-roger
```

```
## 
## Call:
## lm(formula = average_absolute_pointing_error_change ~ spo2 * 
##     intervention, data = hx_dat)
## 
## Residuals:
##     Min      1Q  Median      3Q     Max 
## -70.984  -9.643  -2.049  15.615  42.652 
## 
## Coefficients:
##                               Estimate Std. Error t value Pr(>|t|)  
## (Intercept)                   -45.3497    45.5490  -0.996   0.3213  
## spo2                            0.4872     0.5340   0.912   0.3633  
## interventionintermittent      192.6446    92.2628   2.088   0.0388 *
## spo2:interventionintermittent  -2.1309     1.0594  -2.011   0.0464 *
## ---
## Signif. codes:  0 '***' 0.001 '**' 0.01 '*' 0.05 '.' 0.1 ' ' 1
## 
## Residual standard error: 23.7 on 128 degrees of freedom
## Multiple R-squared:  0.04752,    Adjusted R-squared:  0.02519 
## F-statistic: 2.128 on 3 and 128 DF,  p-value: 0.09982
```

```
##  intervention spo2.trend    SE  df lower.CL upper.CL
##  continuous        0.487 0.534 128   -0.569    1.544
##  intermittent     -1.644 0.915 128   -3.454    0.167
## 
## Confidence level used: 0.95
```

MAPPING R2 FIT

```
## NOTE: Results may be misleading due to involvement in interactions
```

```
##  contrast          estimate   SE  df t.ratio p.value
##  (ses-0) - (ses-1)     27.3 7.29 209   3.747  0.0002
## 
## Results are averaged over the levels of: intervention 
## Degrees-of-freedom method: kenward-roger
```

```
##  contrast                                    estimate    SE    df t.ratio
##  (continuous,ses-0) - (continuous,ses-1)        18.93  7.34 199.2   2.580
##  (continuous,ses-0) - (intermittent,ses-0)     -45.06 17.00  92.8  -2.648
##  (continuous,ses-0) - (intermittent,ses-1)      20.30 17.80 106.0   1.139
##  (continuous,ses-1) - (intermittent,ses-0)     -63.99 17.00  92.0  -3.769
##  (continuous,ses-1) - (intermittent,ses-1)       1.36 17.80 105.1   0.077
##  (intermittent,ses-0) - (intermittent,ses-1)    65.35 12.20 211.9   5.355
##  p.value
##   0.0514
##   0.0462
##   0.6664
##   0.0016
##   0.9998
##   <.0001
## 
## Degrees-of-freedom method: kenward-roger 
## P value adjustment: tukey method for comparing a family of 4 estimates
```

```
## 
## Call:
## lm(formula = mapping_rsquare_change ~ spo2 * intervention, data = hx_dat)
## 
## Residuals:
##      Min       1Q   Median       3Q      Max 
## -0.64917 -0.28099  0.06931  0.18670  0.73111 
## 
## Coefficients:
##                                 Estimate Std. Error t value Pr(>|t|)
## (Intercept)                   -0.0308957  0.5786746  -0.053    0.958
## spo2                          -0.0008673  0.0067842  -0.128    0.898
## interventionintermittent      -0.3697582  1.1721457  -0.315    0.753
## spo2:interventionintermittent  0.0032914  0.0134594   0.245    0.807
## 
## Residual standard error: 0.3011 on 128 degrees of freedom
## Multiple R-squared:  0.01543,    Adjusted R-squared:  -0.007649 
## F-statistic: 0.6685 on 3 and 128 DF,  p-value: 0.5729
```

```
##  intervention spo2.trend      SE  df lower.CL upper.CL
##  continuous    -0.000867 0.00678 128  -0.0143   0.0126
##  intermittent   0.002424 0.01160 128  -0.0206   0.0254
## 
## Confidence level used: 0.95
```

PERSPECTIVE TAKING ERROR

```
## NOTE: Results may be misleading due to involvement in interactions
```

```
##  contrast          estimate   SE  df t.ratio p.value
##  (ses-0) - (ses-1)    -26.4 7.69 214  -3.435  0.0007
## 
## Results are averaged over the levels of: intervention 
## Degrees-of-freedom method: kenward-roger
```

```
##  contrast                                    estimate    SE    df t.ratio
##  (continuous,ses-0) - (continuous,ses-1)       -42.33  7.69 202.8  -5.503
##  (continuous,ses-0) - (intermittent,ses-0)     -31.00 17.30  99.7  -1.789
##  (continuous,ses-0) - (intermittent,ses-1)       2.08 17.80 107.4   0.117
##  (continuous,ses-1) - (intermittent,ses-0)      11.34 17.30  98.8   0.656
##  (continuous,ses-1) - (intermittent,ses-1)      44.41 17.80 106.5   2.497
##  (intermittent,ses-0) - (intermittent,ses-1)    33.07 12.90 218.2   2.569
##  p.value
##   <.0001
##   0.2848
##   0.9994
##   0.9133
##   0.0661
##   0.0527
## 
## Degrees-of-freedom method: kenward-roger 
## P value adjustment: tukey method for comparing a family of 4 estimates
```

```
## 
## Call:
## lm(formula = average_absolute_perspective_taking_error_change ~ 
##     spo2 * intervention, data = hx_dat)
## 
## Residuals:
##     Min      1Q  Median      3Q     Max 
## -39.893  -6.785   2.227   8.080  43.711 
## 
## Coefficients:
##                               Estimate Std. Error t value Pr(>|t|)
## (Intercept)                   -12.1344    29.7154  -0.408    0.684
## spo2                            0.1139     0.3484   0.327    0.744
## interventionintermittent       59.1150    59.9400   0.986    0.326
## spo2:interventionintermittent  -0.7910     0.6889  -1.148    0.253
## 
## Residual standard error: 15.46 on 132 degrees of freedom
## Multiple R-squared:  0.08384,    Adjusted R-squared:  0.06301 
## F-statistic: 4.026 on 3 and 132 DF,  p-value: 0.008882
```

```
##  intervention spo2.trend    SE  df lower.CL upper.CL
##  continuous        0.114 0.348 132   -0.575    0.803
##  intermittent     -0.677 0.594 132   -1.853    0.499
## 
## Confidence level used: 0.95
```

```
## Warning: Removed 6 rows containing missing values or values outside the scale range
## (`position_quasirandom()`).
```

```
## Warning: Removed 6 rows containing missing values or values outside the scale range
## (`geom_line()`).
```

EXPONENT

```
## NOTE: Results may be misleading due to involvement in interactions
```

```
##  contrast          estimate   SE  df t.ratio p.value
##  (ses-0) - (ses-1)     13.9 6.17 106   2.248  0.0267
## 
## Results are averaged over the levels of: intervention, eye_state 
## Degrees-of-freedom method: kenward-roger
```

```
## NOTE: Results may be misleading due to involvement in interactions
```

```
##  contrast                                    estimate    SE    df t.ratio
##  (continuous,ses-0) - (continuous,ses-1)        -3.19  8.33 103.8  -0.383
##  (continuous,ses-0) - (intermittent,ses-0)     -15.89  9.95  68.5  -1.596
##  (continuous,ses-0) - (intermittent,ses-1)      16.89 10.50  74.3   1.614
##  (continuous,ses-1) - (intermittent,ses-0)     -12.70 10.00  67.3  -1.265
##  (continuous,ses-1) - (intermittent,ses-1)      20.07 10.50  73.2   1.904
##  (intermittent,ses-0) - (intermittent,ses-1)    32.77  8.66 108.1   3.784
##  p.value
##   0.9808
##   0.3876
##   0.3770
##   0.5881
##   0.2354
##   0.0014
## 
## Results are averaged over the levels of: eye_state 
## Degrees-of-freedom method: kenward-roger 
## P value adjustment: tukey method for comparing a family of 4 estimates
```

```
## 
## Call:
## lm(formula = exponent_mean_change ~ spo2 * intervention, data = hx_dat)
## 
## Residuals:
##      Min       1Q   Median       3Q      Max 
## -1.12889 -0.10453 -0.02213  0.13058  0.90308 
## 
## Coefficients:
##                                Estimate Std. Error t value Pr(>|t|)
## (Intercept)                    0.403631   1.452842   0.278    0.782
## spo2                          -0.004625   0.016510  -0.280    0.780
## interventionintermittent      -1.216635   1.879283  -0.647    0.520
## spo2:interventionintermittent  0.012040   0.021416   0.562    0.576
## 
## Residual standard error: 0.3282 on 59 degrees of freedom
## Multiple R-squared:  0.06682,    Adjusted R-squared:  0.01937 
## F-statistic: 1.408 on 3 and 59 DF,  p-value: 0.2493
```

```
##  intervention spo2.trend     SE df lower.CL upper.CL
##  continuous     -0.00463 0.0165 59  -0.0377   0.0284
##  intermittent    0.00741 0.0136 59  -0.0199   0.0347
## 
## Confidence level used: 0.95
```

OFFSET

```
## 
## Call:
## lm(formula = offset_mean_change ~ spo2 * intervention, data = hx_dat)
## 
## Residuals:
##     Min      1Q  Median      3Q     Max 
## -5.0668 -0.0919  0.0562  0.3197  0.9536 
## 
## Coefficients:
##                               Estimate Std. Error t value Pr(>|t|)
## (Intercept)                    4.35701    3.77538   1.154    0.253
## spo2                          -0.04969    0.04290  -1.158    0.252
## interventionintermittent      -5.41279    4.88354  -1.108    0.272
## spo2:interventionintermittent  0.05914    0.05565   1.063    0.292
## 
## Residual standard error: 0.8527 on 59 degrees of freedom
## Multiple R-squared:  0.03969,    Adjusted R-squared:  -0.009136 
## F-statistic: 0.8129 on 3 and 59 DF,  p-value: 0.4918
```

```
##  intervention spo2.trend     SE df lower.CL upper.CL
##  continuous     -0.04969 0.0429 59  -0.1355   0.0362
##  intermittent    0.00946 0.0354 59  -0.0615   0.0804
## 
## Confidence level used: 0.95
```

ALPHA POWER

```
## NOTE: Results may be misleading due to involvement in interactions
```

```
##  contrast                  estimate   SE   df t.ratio p.value
##  continuous - intermittent     16.8 7.91 36.5   2.128  0.0401
## 
## Results are averaged over the levels of: session, eye_state 
## Degrees-of-freedom method: kenward-roger
```

```
## NOTE: Results may be misleading due to involvement in interactions
```

```
##  contrast          estimate   SE   df t.ratio p.value
##  (ses-0) - (ses-1)     19.1 5.79 91.3   3.293  0.0014
## 
## Results are averaged over the levels of: intervention, eye_state 
## Degrees-of-freedom method: kenward-roger
```

```
## NOTE: Results may be misleading due to involvement in interactions
```

```
##  contrast                          estimate    SE   df t.ratio p.value
##  continuous,EC - continuous,EO       -16.77  7.73 80.7  -2.169  0.1406
##  continuous,EC - intermittent,EC       5.84  9.21 66.2   0.633  0.9209
##  continuous,EC - intermittent,EO      11.58  9.79 72.9   1.183  0.6397
##  continuous,EO - intermittent,EC      22.61  9.64 70.7   2.345  0.0975
##  continuous,EO - intermittent,EO      28.35 10.20 76.8   2.782  0.0336
##  intermittent,EC - intermittent,EO     5.74  8.30 83.7   0.691  0.9002
## 
## Results are averaged over the levels of: session 
## Degrees-of-freedom method: kenward-roger 
## P value adjustment: tukey method for comparing a family of 4 estimates
```

```
## 
## Call:
## lm(formula = alpha_pow_mean_change ~ spo2 * intervention, data = hx_dat)
## 
## Residuals:
##     Min      1Q  Median      3Q     Max 
## -0.7472 -0.1424 -0.0425  0.1282  0.7127 
## 
## Coefficients:
##                               Estimate Std. Error t value Pr(>|t|)  
## (Intercept)                    1.06275    1.26953   0.837   0.4066  
## spo2                          -0.01147    0.01439  -0.797   0.4294  
## interventionintermittent      -2.82201    1.65915  -1.701   0.0953 .
## spo2:interventionintermittent  0.03108    0.01882   1.652   0.1050  
## ---
## Signif. codes:  0 '***' 0.001 '**' 0.01 '*' 0.05 '.' 0.1 ' ' 1
## 
## Residual standard error: 0.2596 on 49 degrees of freedom
## Multiple R-squared:  0.08656,    Adjusted R-squared:  0.03063 
## F-statistic: 1.548 on 3 and 49 DF,  p-value: 0.214
```

```
##  intervention spo2.trend     SE df lower.CL upper.CL
##  continuous      -0.0115 0.0144 49 -0.04039   0.0175
##  intermittent     0.0196 0.0121 49 -0.00474   0.0440
## 
## Confidence level used: 0.95
```

ALPHA FREQUENCY

```
## 
## Call:
## lm(formula = alpha_freq_mean_change ~ spo2 * intervention, data = hx_dat)
## 
## Residuals:
##      Min       1Q   Median       3Q      Max 
## -1.73460 -0.20180  0.01439  0.14564  1.32997 
## 
## Coefficients:
##                               Estimate Std. Error t value Pr(>|t|)
## (Intercept)                   -2.74639    2.31067  -1.189    0.240
## spo2                           0.03088    0.02619   1.179    0.244
## interventionintermittent       1.82913    3.01981   0.606    0.548
## spo2:interventionintermittent -0.02205    0.03425  -0.644    0.523
## 
## Residual standard error: 0.4726 on 49 degrees of freedom
## Multiple R-squared:  0.04551,    Adjusted R-squared:  -0.01293 
## F-statistic: 0.7787 on 3 and 49 DF,  p-value: 0.5115
```

BETA POWER

```
## NOTE: Results may be misleading due to involvement in interactions
```

```
##  contrast                  estimate   SE   df t.ratio p.value
##  continuous - intermittent     16.2 8.27 33.5   1.954  0.0590
## 
## Results are averaged over the levels of: session, eye_state 
## Degrees-of-freedom method: kenward-roger
```

```
## NOTE: Results may be misleading due to involvement in interactions
```

```
##  contrast          estimate   SE   df t.ratio p.value
##  (ses-0) - (ses-1)     25.7 5.06 83.3   5.068  <.0001
## 
## Results are averaged over the levels of: intervention, eye_state 
## Degrees-of-freedom method: kenward-roger
```

```
## NOTE: Results may be misleading due to involvement in interactions
```

```
##  contrast                          estimate   SE   df t.ratio p.value
##  continuous,EC - continuous,EO        -8.65 6.68 71.2  -1.296  0.5686
##  continuous,EC - intermittent,EC       5.86 9.60 55.9   0.610  0.9284
##  continuous,EC - intermittent,EO      18.30 9.67 58.2   1.892  0.2428
##  continuous,EO - intermittent,EC      14.51 9.56 54.9   1.517  0.4342
##  continuous,EO - intermittent,EO      26.95 9.64 57.3   2.796  0.0344
##  intermittent,EC - intermittent,EO    12.44 7.17 75.2   1.735  0.3130
## 
## Results are averaged over the levels of: session 
## Degrees-of-freedom method: kenward-roger 
## P value adjustment: tukey method for comparing a family of 4 estimates
```

```
## 
## Call:
## lm(formula = beta_pow_mean_change ~ spo2 * intervention, data = hx_dat)
## 
## Residuals:
##      Min       1Q   Median       3Q      Max 
## -0.16100 -0.06534 -0.01604  0.06029  0.35096 
## 
## Coefficients:
##                                 Estimate Std. Error t value Pr(>|t|)
## (Intercept)                    0.0751424  0.5279730   0.142    0.887
## spo2                          -0.0008322  0.0059759  -0.139    0.890
## interventionintermittent       0.0989311  0.6685884   0.148    0.883
## spo2:interventionintermittent -0.0017046  0.0076389  -0.223    0.824
## 
## Residual standard error: 0.09922 on 47 degrees of freedom
## Multiple R-squared:  0.05983,    Adjusted R-squared:  -0.0001824 
## F-statistic: 0.997 on 3 and 47 DF,  p-value: 0.4025
```

BETA FREQUENCY

```
## boundary (singular) fit: see help('isSingular')
```

```
## NOTE: Results may be misleading due to involvement in interactions
```

```
##  contrast                  estimate   SE   df t.ratio p.value
##  continuous - intermittent     12.5 6.44 30.3   1.940  0.0617
## 
## Results are averaged over the levels of: session, eye_state 
## Degrees-of-freedom method: kenward-roger
```

```
## boundary (singular) fit: see help('isSingular')
```

```
## NOTE: Results may be misleading due to involvement in interactions
```

```
##  contrast          estimate   SE   df t.ratio p.value
##  (ses-0) - (ses-1)    -13.1 6.23 89.6  -2.096  0.0389
## 
## Results are averaged over the levels of: intervention, eye_state 
## Degrees-of-freedom method: kenward-roger
```

```
## 
## Call:
## lm(formula = beta_freq_mean_change ~ spo2 * intervention, data = hx_dat)
## 
## Residuals:
##     Min      1Q  Median      3Q     Max 
## -6.5100 -0.8594  0.1238  0.9873  5.1226 
## 
## Coefficients:
##                               Estimate Std. Error t value Pr(>|t|)
## (Intercept)                    13.2943    11.8989   1.117    0.270
## spo2                           -0.1475     0.1347  -1.095    0.279
## interventionintermittent        0.9426    15.0680   0.063    0.950
## spo2:interventionintermittent  -0.0172     0.1722  -0.100    0.921
## 
## Residual standard error: 2.236 on 47 degrees of freedom
## Multiple R-squared:  0.07243,    Adjusted R-squared:  0.01322 
## F-statistic: 1.223 on 3 and 47 DF,  p-value: 0.3117
```

```
## Warning: package 'knitr' was built under R version 4.5.3
```

```
## Warning: package 'kableExtra' was built under R version 4.5.3
```

```
## 
## Attaching package: 'kableExtra'
```

```
## The following object is masked from 'package:dplyr':
## 
##     group_rows
```
